# Background free tracking of single RNA in living cells using catalytically inactive *Cas*E

**DOI:** 10.1101/635912

**Authors:** Feng Gao, Yue Sun, Feng Jiang, Xiaoyue Bai, Chunyu Han

**Affiliations:** Gene Editing Research Center, Hebei University of Science and Technology, Shijiazhuang, Hebei 050018, China

## Abstract

RNAs have important and diverse functions. Visualizing an isolated RNA in living cells provide us essential information of its roles. By now, there are two kinds of live RNA imaging systems invented, one is the MS2 system and the other is the *Cas*13a system. In this study, we show that when fused with split-Fp, *Cas*E can be engineered into a live RNA tracking tool.

## Introduction

*Cas*E is a core component of type I-E CRISPR complex, which solely processes pre-crRNA by binding specific stem-loop region, which we called *<u>C</u>as*E <u>B</u>inding <u>S</u>ite (CBS). Even with restriction digest, it still tightly binds to the 3’-terminus of mature crRNA^1^ (Figure 1A). The conserved His^20^ of *Cas*E is involved in the catalysis activity^2^; The identical mutation, “ΔHis26-T*tCse3*, lost the catalytic activity but still tightly binds to its target^3^. In this study, by conjugating the split Venus to the N-and C-terminus of d*Cas*E (“ΔHis20), a live RNA tracking tool, VN-d*Cas*E-VC, was constructed. This system emits florescence only when the target RNAs are present, thus enhancing the signal to noise performance.

**Figure 1A.**
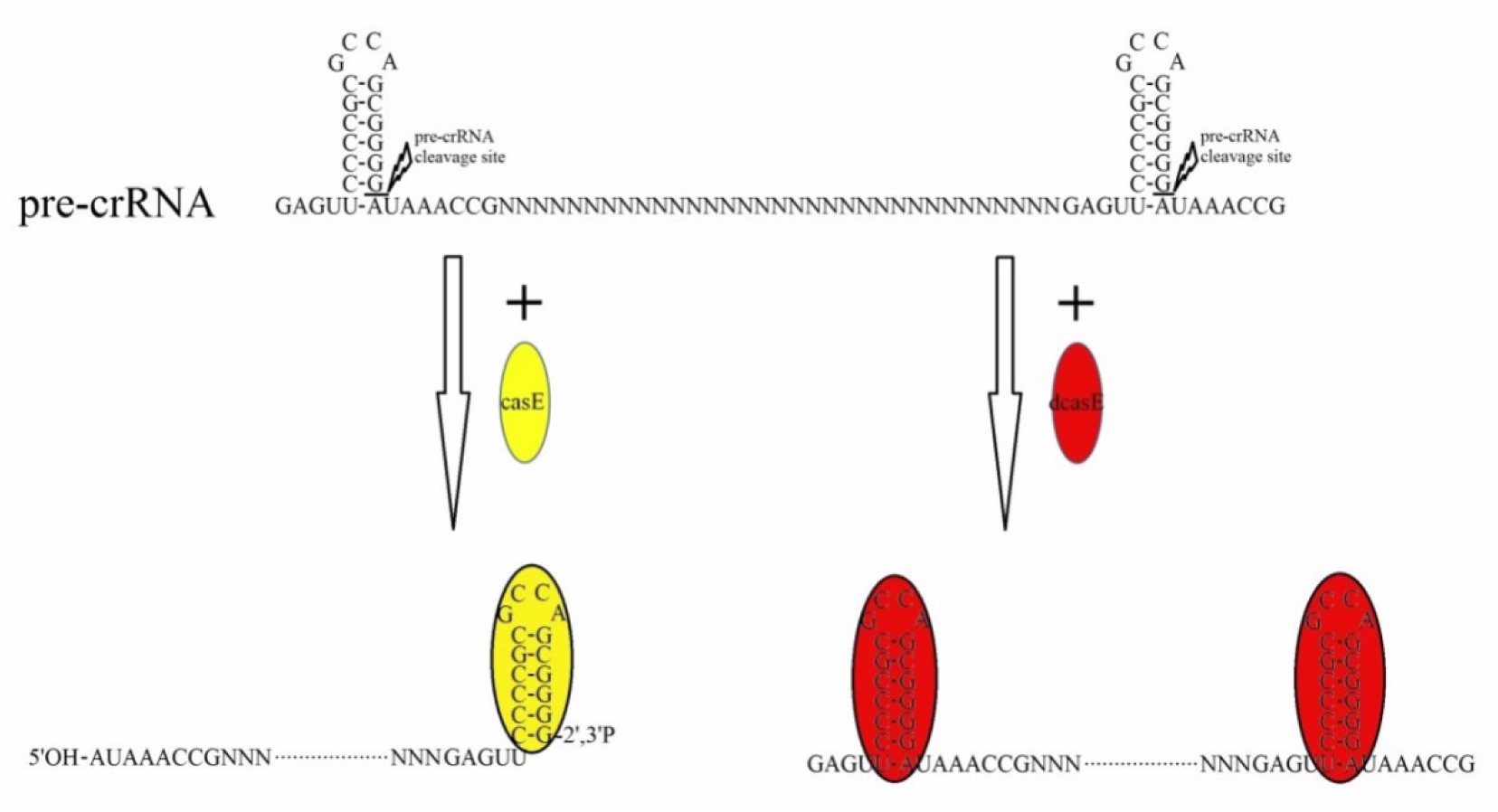
Graphical illustration of the *Cas*E and d*Cas*E. *Cas*E binds and cleaves the stem-loop handle (CBS) of target RNA, while d*Cas*E only binds the CBS^1,2,3^.

## Results

Visualizing RNA in living cells requires the tracking tool without preference for specific sub-cellular distribution. The high-level expression and even distribution of *Cas*E-GFP and d*Cas*E-GFP in HEK293T cells indicated that *Cas*E and d*Cas*E are suitable for RNA manipulation in living cells (Figure 1B).

**Figure 1B.**
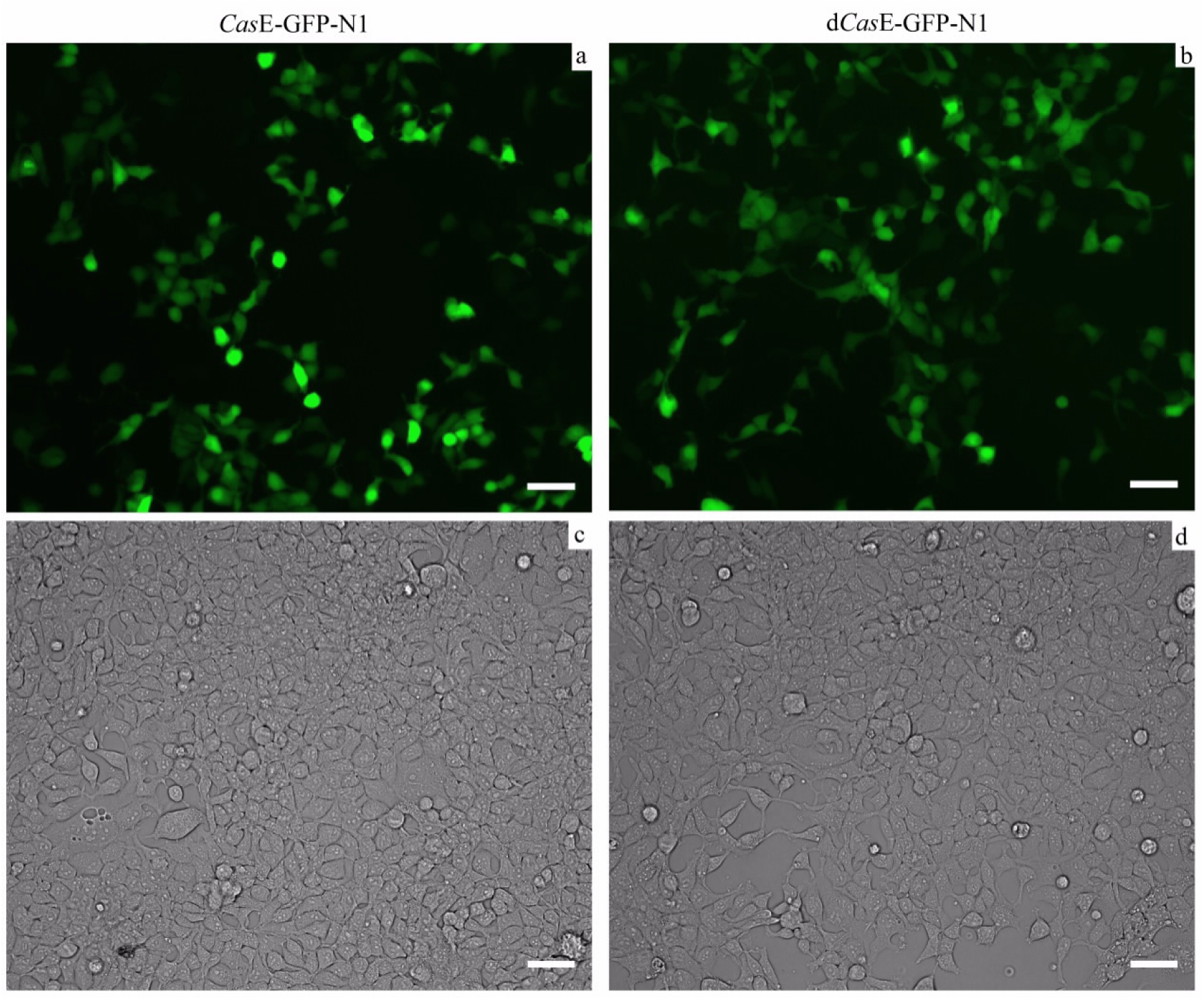
Overexpression of *Cas*E-GFP and d*Cas*E-GFP by transient transfection of 293T cells cultivated in a 24-well plate (2 cm^2^). The transfection dosage is 300 ng plasmid per well. c and d are the bright field images of a and b. Scale bar indicates 50 μm.

To test whether *Cas*E has robust activity when expressed in mammalian cells, a “turn-off” reporter, plasmid CBS-GFP-N1, was constructed. It consisted of a GFP mRNA and a *<u>C</u>as*E <u>B</u>inding <u>S</u>ite (CBS) in its 5’-UTR. When *Cas*E was introduced into the system, the GFP expression level was sharply reduced (Figures 1C and 1D). To further test the *Cas*E activity, a “turn-on” reporter, plasmid RED-16×CBS-Lin28-C1, was also constructed, in which the CBS was inserted in the 3’-UTR region of RED monomer gene and upstream of Lin28. Lin28 is an RNA nuclear retain signal, the RED monomer mRNA with a lin28 signal in its 3’-UTR can hardly be translated into protein^4^. The CBS-*Cas*E dependent restriction cuts off the lin28 signal and release the target mRNA from nuclear for translation (Figure 1E and Figure 1F). These indicated that *Cas*E can bind and cleave the CBS in mammalian cells.

**Figure 1C.**
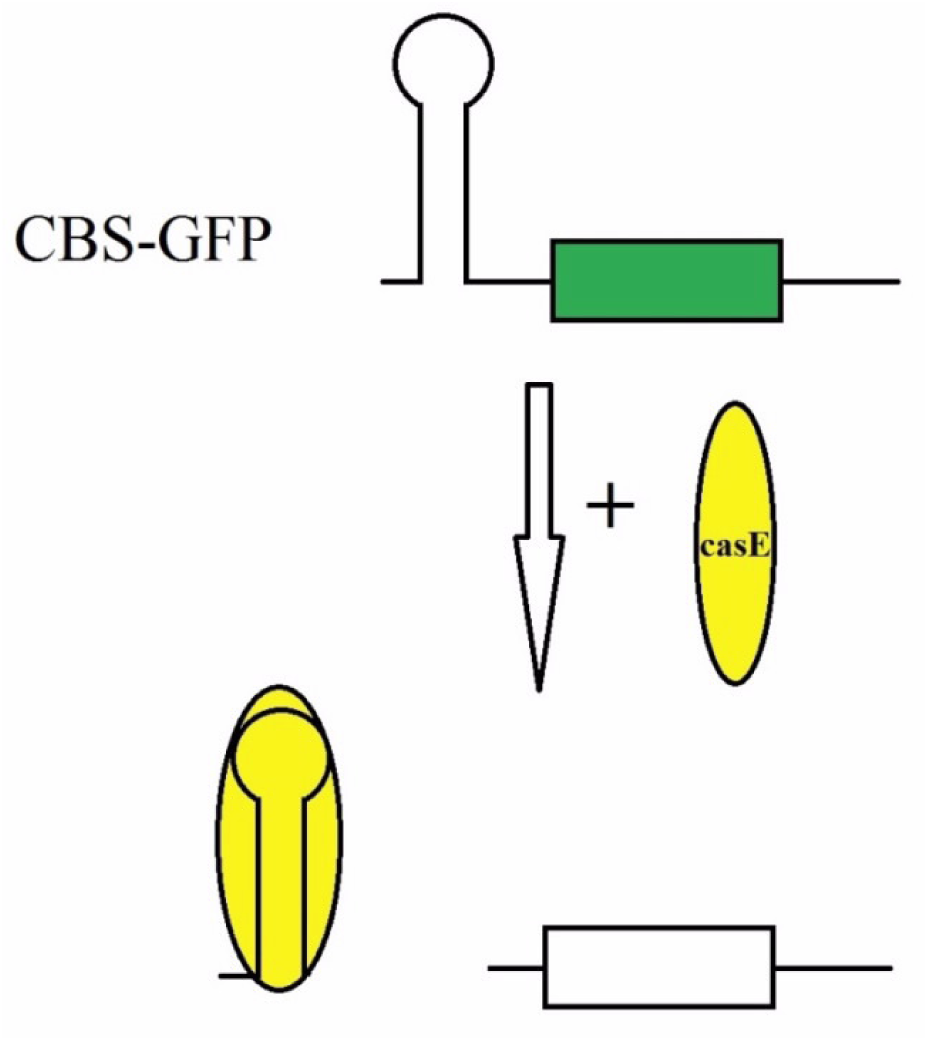
Graphical illustration of the ‘turn-off’ reporter system. *Cas*E binds the CBS region and cleaves the CBS-GFP mRNA. GFP translation was then blocked by decapitation of the 5’-UTR.

**Figure 1D.**
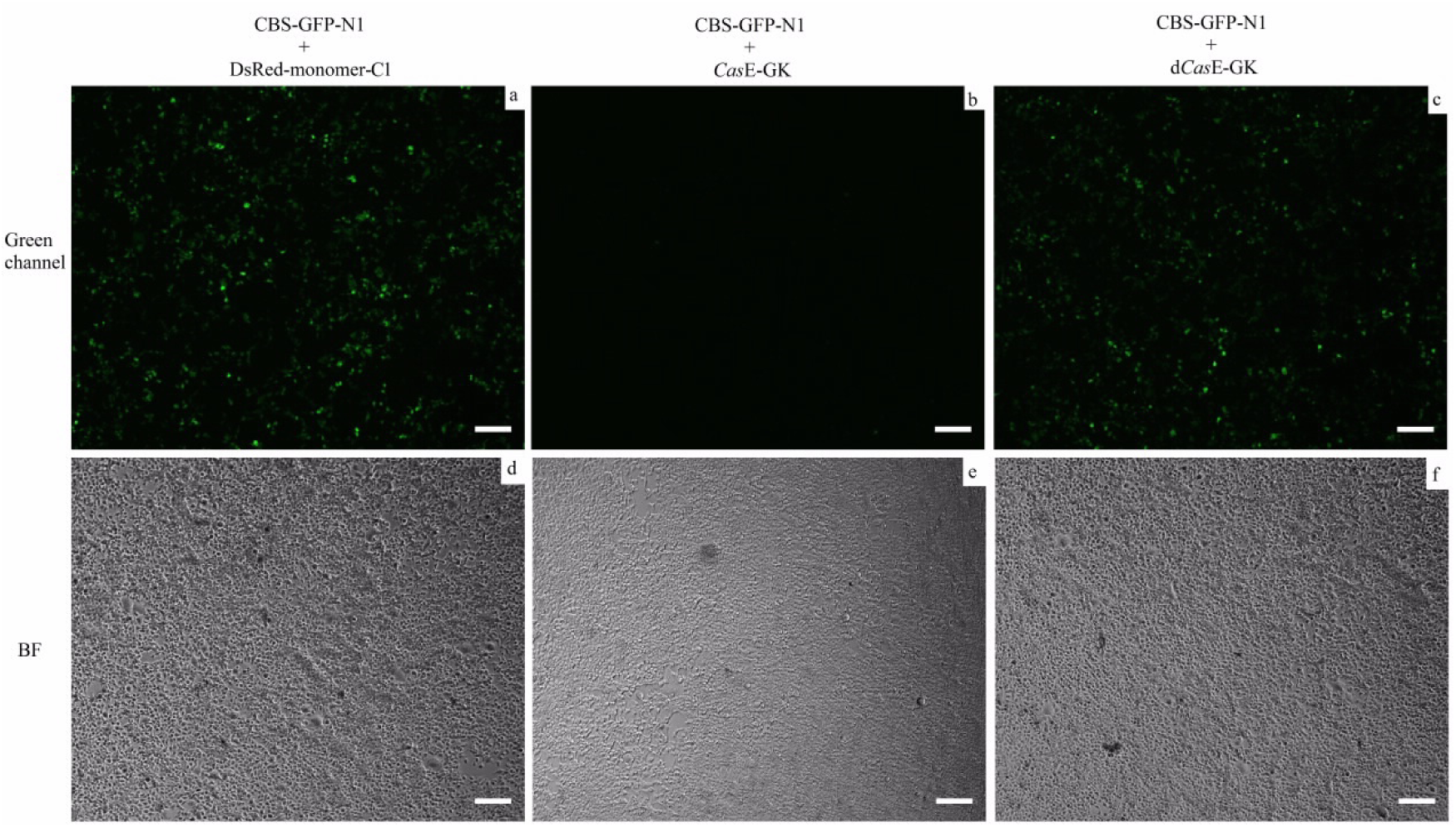
CBS-GFP expression level is sharply decreased only when *Cas*E protein is introduced into the system, but not RFP (negative control) or d*Cas*E. Plasmids combinations for each transfection are indicated on top of the pictures. Plasmids dosages (for one well of a 24-well plate) are: 300 ng of CBS-GFP-N1 and 100 ng of DsRed-monomer-C1, *Cas*E-GK, or d*Cas*E-GK. d–f are bright field images of a–c. Scale bar, 200 μm.

**Figure 1E.**
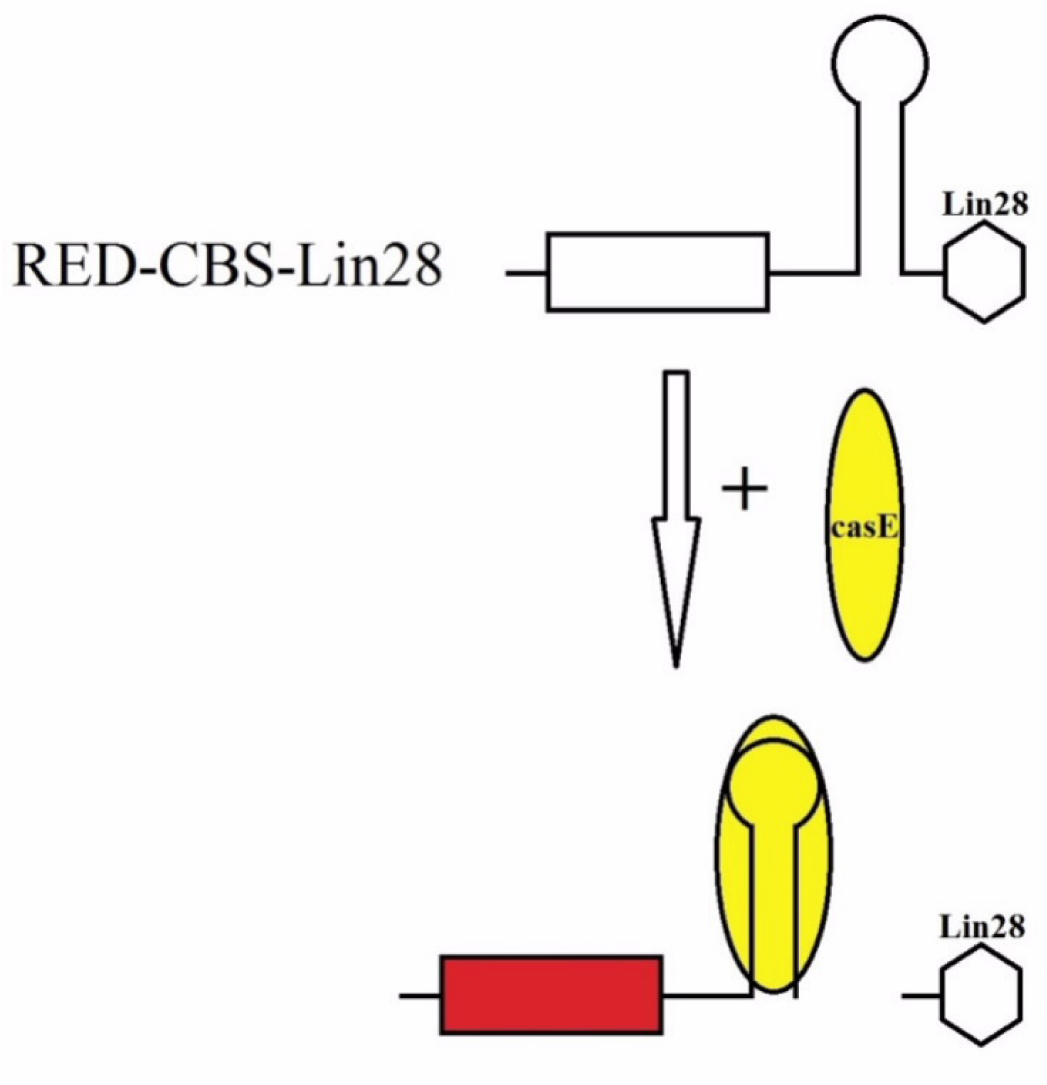
Graphical illustration of the’turn-on’ reporter system. Lin28 is cleaved by *Cas*E, and the RED-CBS mRNA is then released from cell nucleus to cytoplasm for translation.

**Figure 1F.**
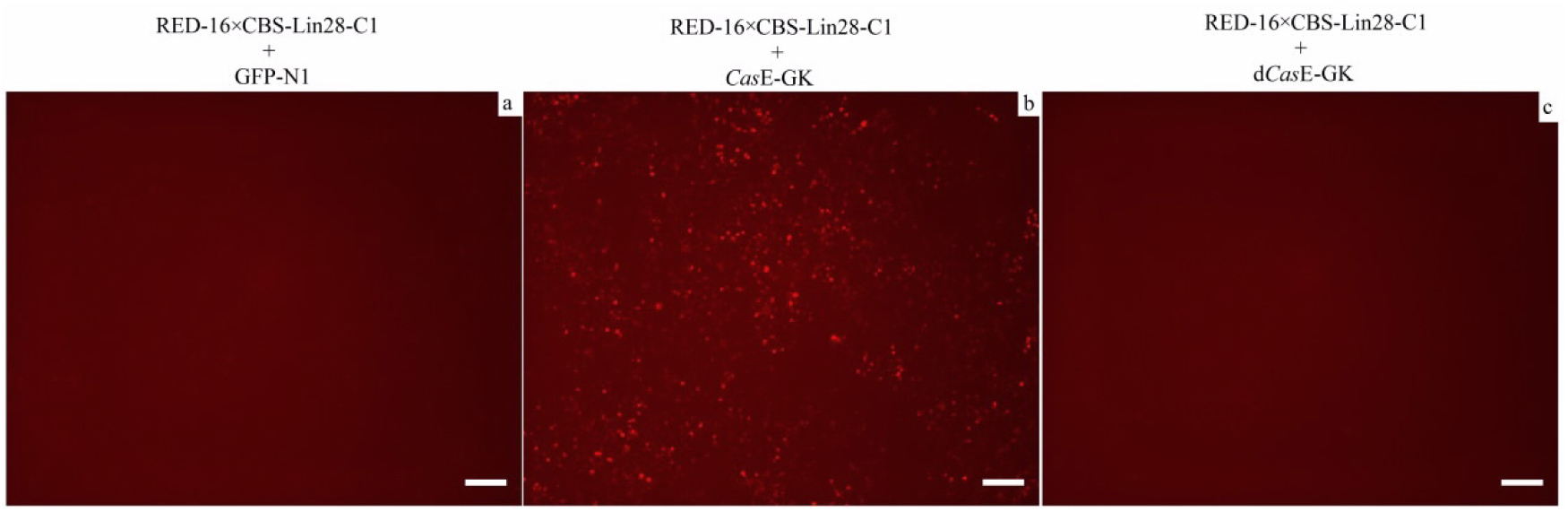
RED-CBS-Lin28 reporter is turned on (red fluorescent) by *Cas*E restriction but not EGFP (negative control) or d*Cas*E protein. The transfection plasmid dosage (for one well of a 24-well plate) are 100ng of RED-16×CBS-Lin28-C1 and 300ng of GFP-N1, *Cas*E-GK or d*Cas*E-GK. Scale bar, 200 μm.

Aiming at engineering the d*Cas*E-CBS interaction into a RNA tracking system, we tried many schemes by conjugating the split-FP^5–7^ to the d*Cas*E protein. We found that one version, the VN-d*Cas*E-VC can hardly emit fluorescent which may due to un-properly folding, or un-stable state^3^, but when the target RNA (CBS) was bound, the Venus signal can be clearly captured under fluorescent microscopy even when the transfection dosage of VN-d*Cas*E-VC expression plasmid is very low (Figure 2A and Figure 2B).

**Figure 2A.**
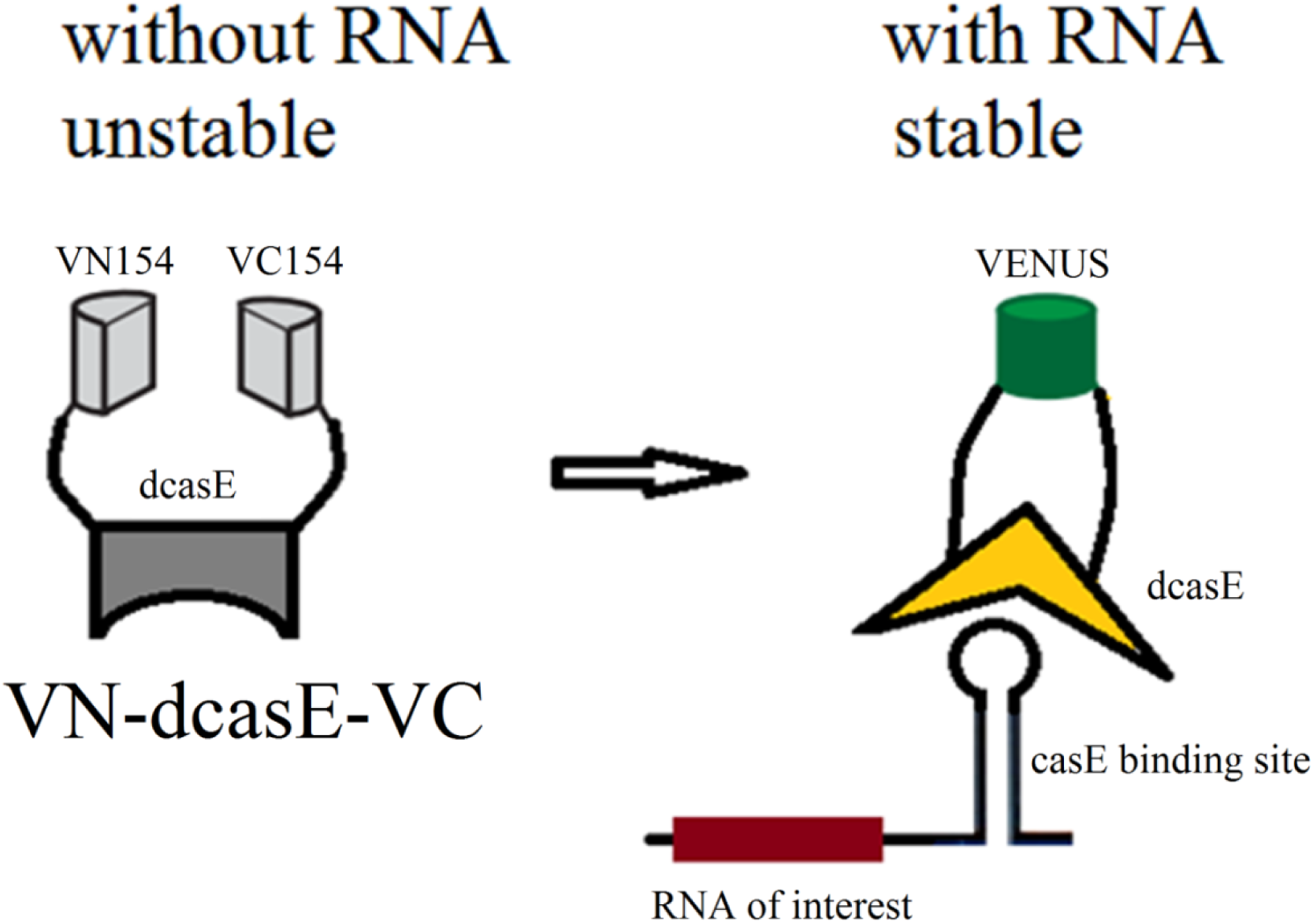
conjectural mechanism for the target induced fluorescent activation of VN-d*Cas*E-VC.

**Figure 2B.**
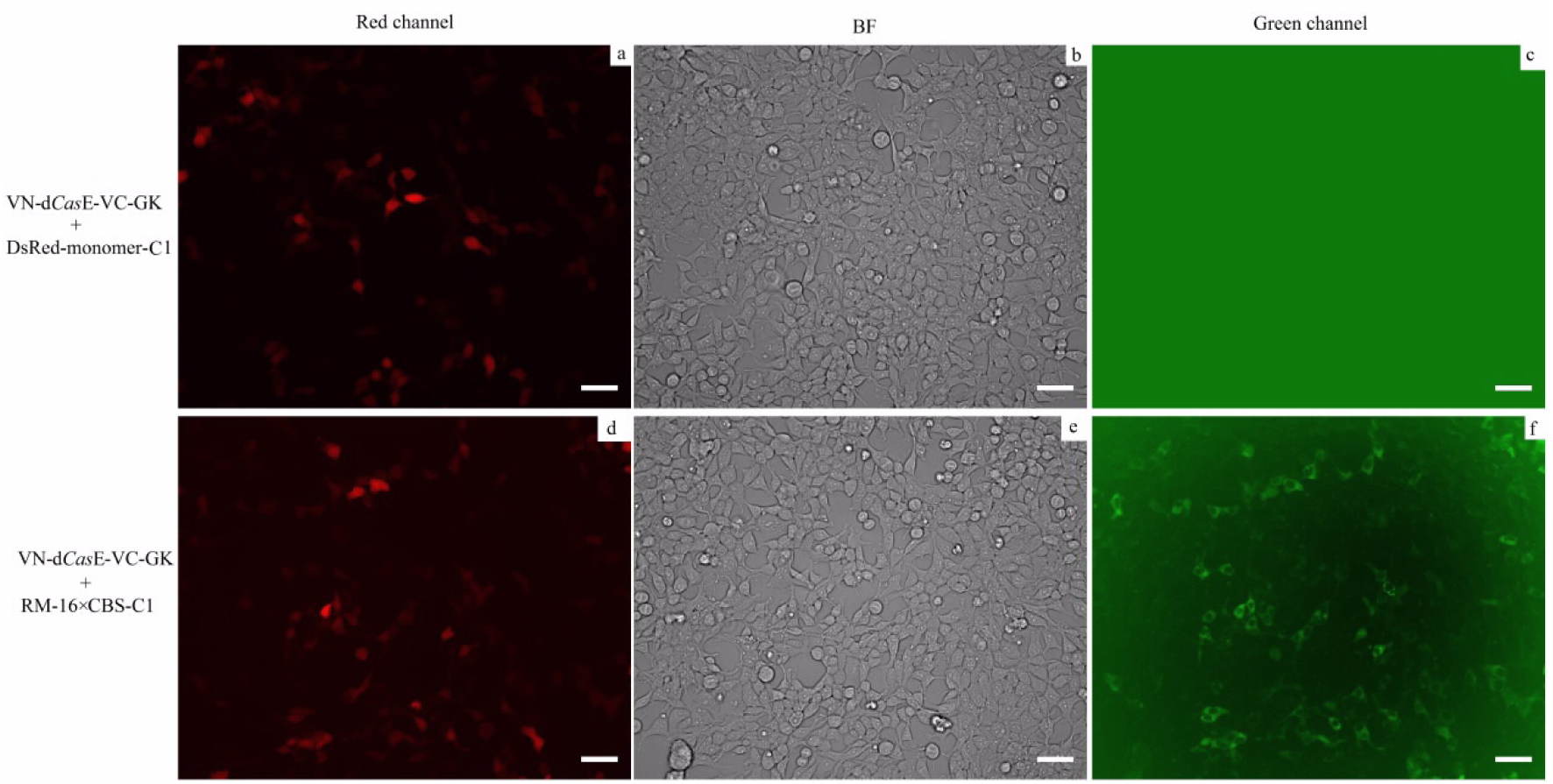
Fluorescent of the VN-d*Cas*E-VC induced by target mRNA in 293T cells. Plasmid RM-16×CBS-C1 derived from vector DsRed-monomer-C1 contains 16×CBS RNAs at the 3’-UTR region of RED monomer gene. VN-d*Cas*E-VC has no fluorescent signal without CBS (c), and induced signal is detected when CBS (target mRNA) is present (f). The transfection plasmid dosages are 50 ng of VN-d*Cas*E-VC-GK and 800 ng of DsRed-monomer-C1 or RM-16×CBS-C1. Scale bar, 50 μm.

The VN-d*Cas*E-VC system was then used to track the overexpressed β-actin mRNA in mammalian cells, with the MS2 system^8–11^ as a control. As shown in Figure 2C, both systems indicated the same localization of β-actin mRNA. However, the MS2 system showed vague image. It might be ascribed to excessive <u>M</u>S2 <u>C</u>oat <u>P</u>rotein (MCP) without target mRNA binding. In contrast, the VN-d*Cas*E-VC system showed strong fluorescence in the cytoplasm (Figure 2C).

**Figure 2C.**
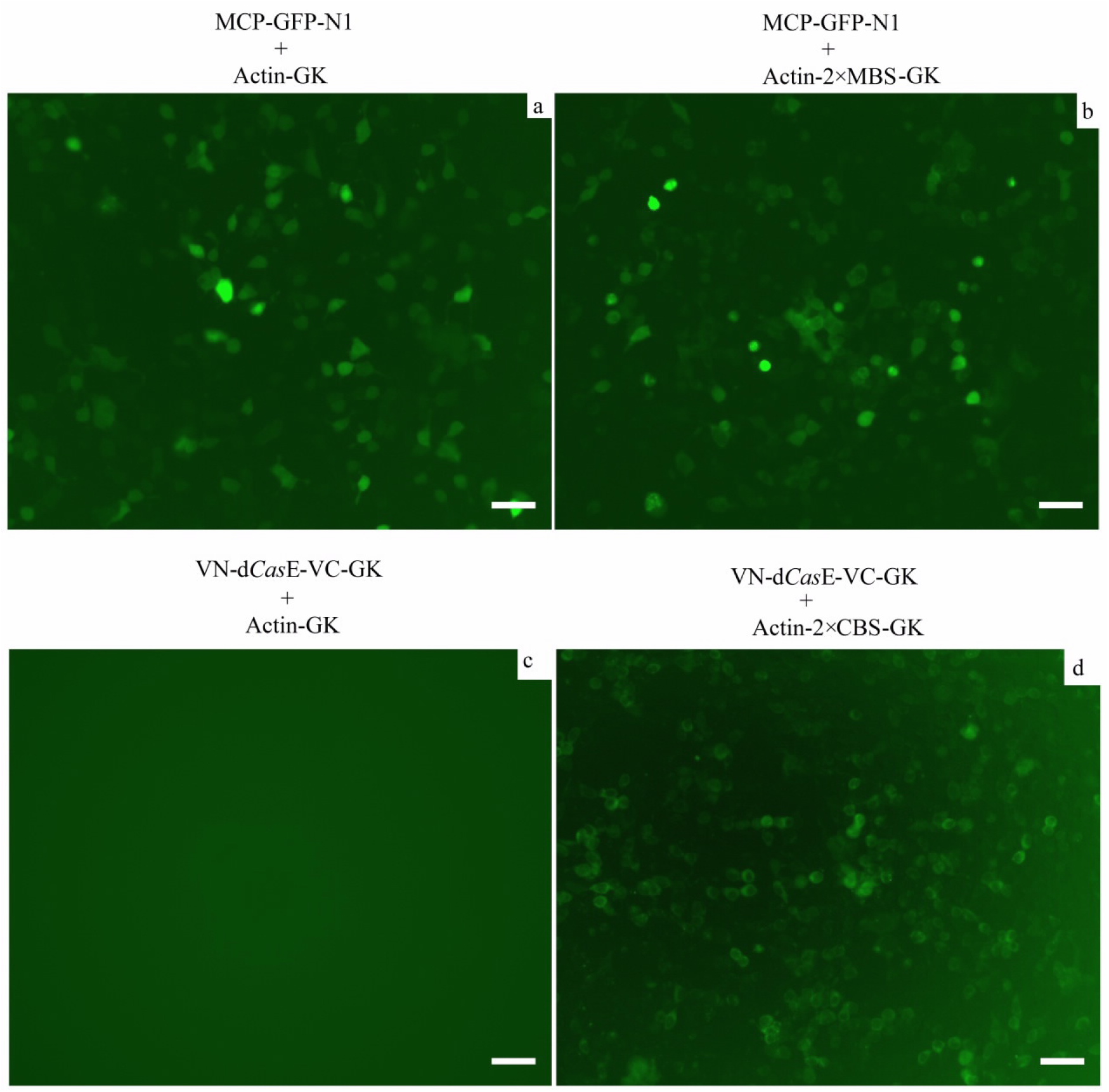
β-Actin mRNA imaging by the VN-d*Cas*E-VC and MS2 systems. Plasmids MCP-GFP-N1 and VN-d*Cas*E-VC-GK are used to overexpress MCP-GFP and VN-d*Cas*E-VC, respectively. Actin-GK, Actin-2×MBS-GK, and Actin-2×CBS-GK are used to transcribe β-actin mRNA without tag, with two <u>M</u>CP <u>B</u>inding <u>S</u>ite (MBS) in the 3’-UTR region, and with two CBS RNAs in the 3’-UTR region, respectively. The transfection plasmid dosages are 50 ng of MCP-GFP-N1 or VN-d*Cas*E-VC-GK, and 800 ng of Actin-GK, Actin-2×MBS-GK, or Actin-2×CBS-GK. Scale bar, 50 μm.

We also observed that, adding more CBS to the target mRNA improves signal, although 2×CBS per molecule is enough for imaging the overexpressed β-actin mRNA (Figure 2D). Thus low abundant mRNA can be detected by increasing the number of CBS to compensate the low concentration.

**Figure 2D.**
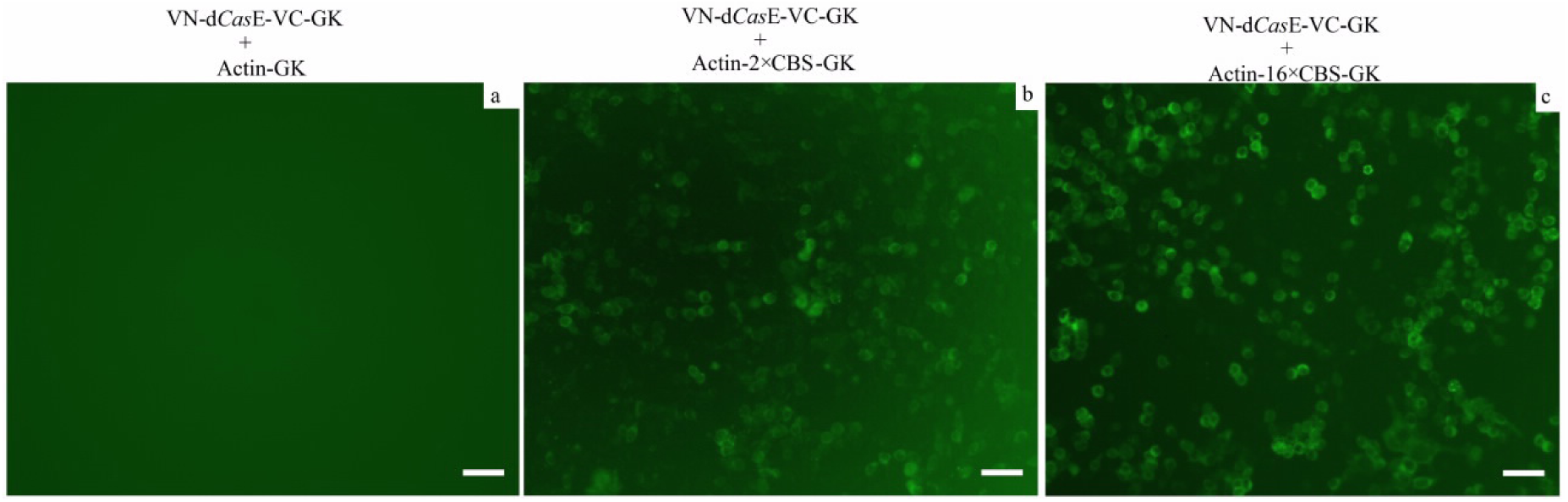
β-Actin mRNA imaging with 0, 2 or 16 CBSs. The transfection plasmid dosages are 50 ng of VN-d*Cas*E-VC-GK and 800 ng of Actin-GK or Actin-2(16)×CBS-GK. Scale bar, 50 μm.

Applying the VN-d*Cas*E-VC system in imaging RNAs displays some interestingresults: 2×CBS RNA transcribed from H1 promoter mainly distributed in nuclear, while 16×CBS RNA from CMV promoter mainly located in cytoplasm(Figure 2E). This preliminary result may reveal that the length of RNA or type of promoter might affect the subcellular localization of RNA.

**Figure 2E.**
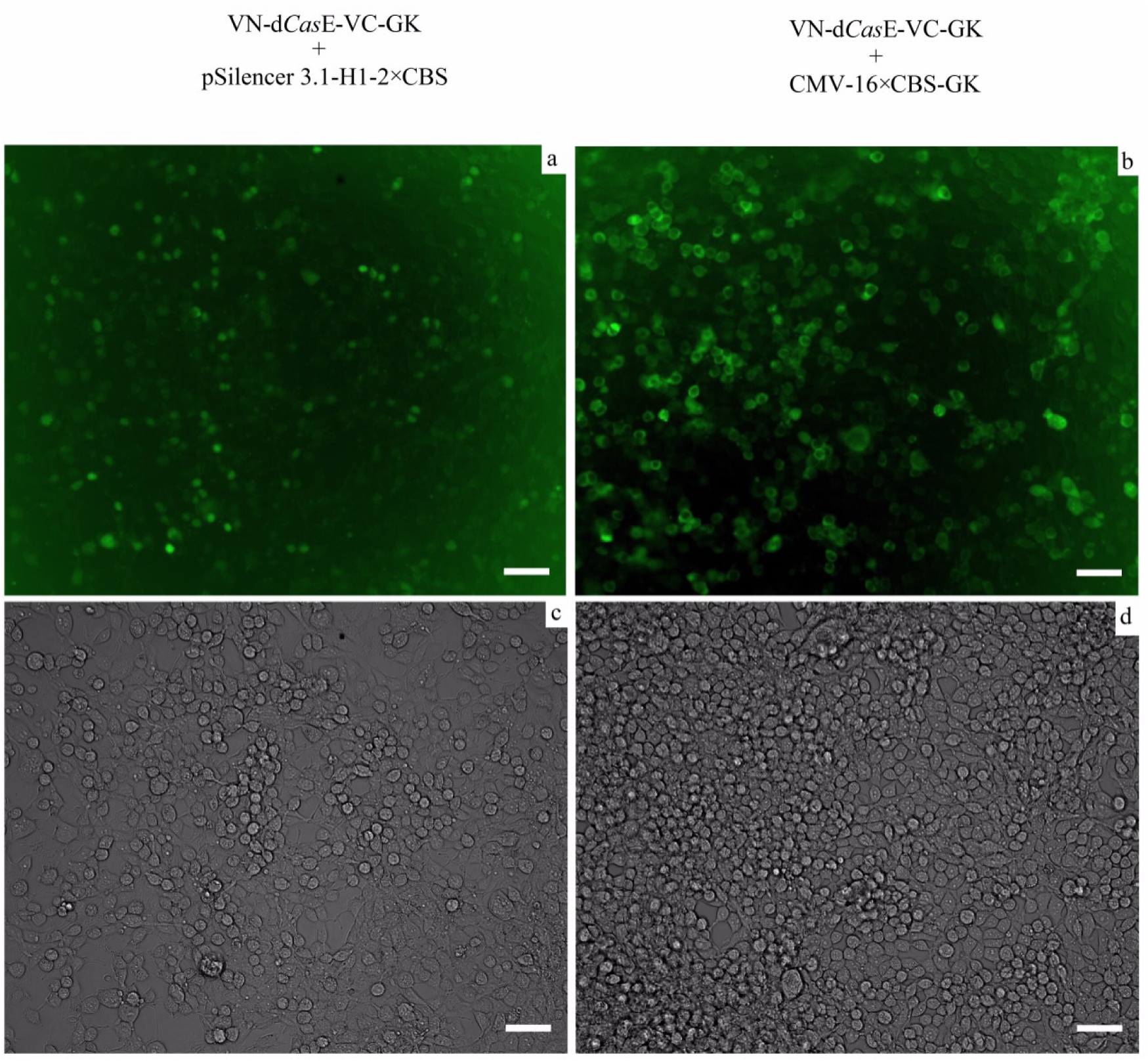
Images of CBS RNAs transcribed from different promoters. Plasmid pSilencer 3.1-H1-2×CBS transcribes 2×CBS RNAs from H1 promoter, while plasmid CMV-16×CBS-GK transcribes 16×CBS RNAs from CMV promoter. c and d are bright fieds of a and b, respectively. The transfection plasmid dosages are 50 ng of VN-d*Cas*E-VC-GK and 800 ng of pSilencer 3.1-H1-2×CBS or CMV-16×CBS-GK. Scale bar, 50 μm.

## Discussion

We invented an new RNA tracking tool, which named VN-dCasE-VC. This system is able to tracking specific RNA without background in living cells. Small Molecular Weight of d*Cas*E make it fit for live RNA tracking: d*Cas*E is only a little bit bigger than MCP but much smaller than *Cas*13a (13 kDa of MCP; 22 kDa of dCasE; more than 130 kDa of *Cas*13a). More than that, VN-d*Cas*E-VC system needs much less target regions than that of MS2 system: 2×CBS is enough for VN-d*Cas*E-VC to obtain high resolution image (Figure 2C) while at least 24×<u>M</u>CP <u>B</u>inding <u>S</u>ite (MBS) is needed for MS2 system^9, 11^. To further improve the performance of VN-d*Cas*E-VC system, optimization of the fluorescent protein segmentation or screening of more Cas orthologues will be applied.

