## supplemental for "Background free tracking of single RNA in living cells using catalytically inactive *Cas*E"

Supplementary data includes methods, 16 constructs, and key sequences.

**Supplementary Figure S1.** *Cas*E-GFP-N1 construct.

**Supplementary Figure S2.** d*Cas*E-GFP-N1 construct.

**Supplementary Figure S3.** MCP-GFP-N1 construct.

**Supplementary Figure S4.** *Cas*E-GK construct.

**Supplementary Figure S5.** d*Cas*E-GK construct.

**Supplementary Figure S6.** VN-d*Cas*E-VC-GK construct.

**Supplementary Figure S7.** RM-16×CBS-C1 construct.

**Supplementary Figure S8.** Actin-GK construct.

**Supplementary Figure S9.** Actin-2×CBS-GK construct.

**Supplementary Figure S10.** Actin-16×CBS-GK construct.

**Supplementary Figure S11.** Actin-2×MBS-GK construct.

**Supplementary Figure S12.** CBS-GFP-N1 construct.

**Supplementary Figure S13.** RED-16×CBS-Lin28-C1 construct.

**Supplementary Figure S14.** pSilencer 3.1-H1-2×CBS construct.

**Supplementary Figure S15.** CMV-16×CBS-GK construct.

**Supplementary Figure S16.** GK construct.

**Methods**

Cell culture and transfection

HEK293T cells (ATCC, CRL-11268) were maintained in Dulbecco’s modified Eagle medium (Gibco, 11965-092) supplemented with 10% fetal bovine serum (Sigma-Aldrich, F4135) and 1% penicillin/streptomycin (Invitrogen, 15140-122). Transient transfection was performed using the Entranster^TM^ R-4000 transfection reagent (Engreen Biosystem, 4000-4) according to the manufacturer’s instruction.

**Supplementary Figure S1**

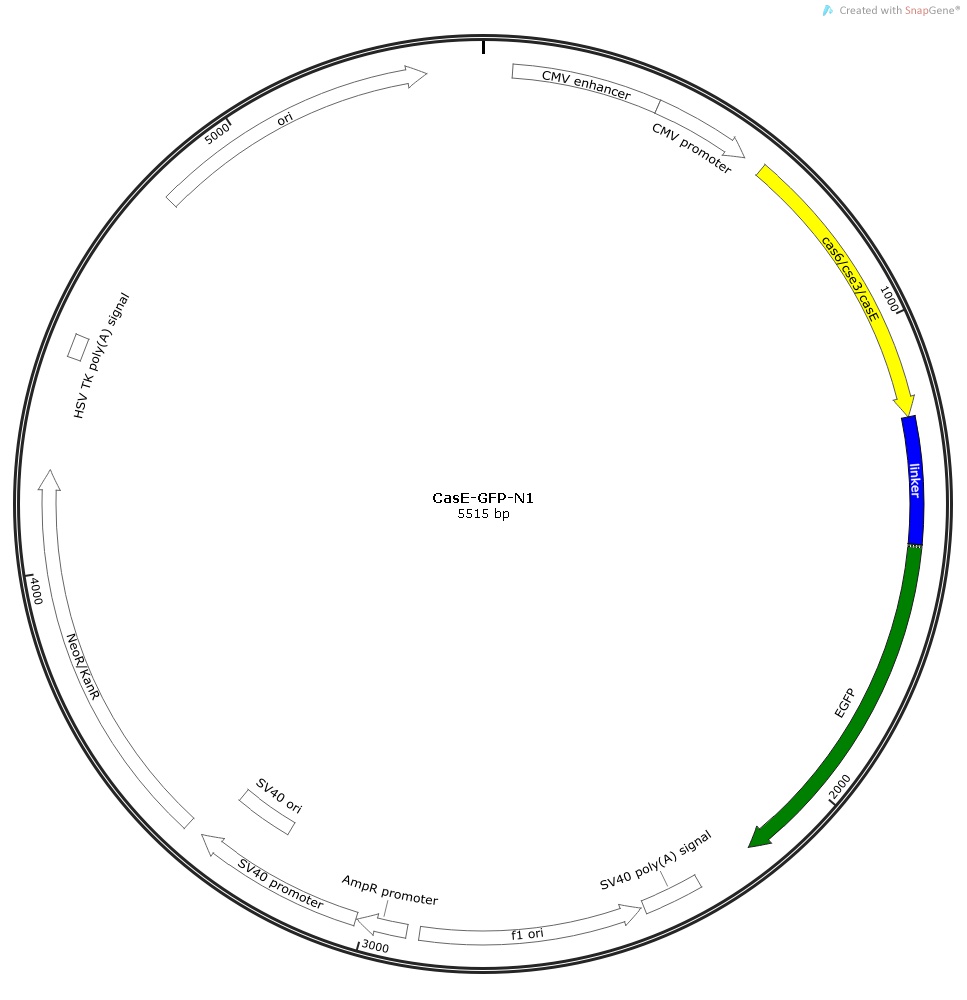

**Supplementary Figure S1.** *Cas*E-GFP-N1 construct.

The coding sequences of *Cas*E, linker, and GFP are shown in yellow, blue, and green, respectively.

ATGGTGtatctcagtaaagtcatcattgccagggcctggagcagggatctttaccaacttcaccagggattatggcatttatttccaaacagaccggatgctgctcgtgattttctttttcatgttgagaagcgaaacacaccagaaggctgtcatgttttattgcagtcagcgcaaatgcctgtttcaactgccgttgcgacagtcattaaaactaaacaggttgaatttcaacttcaggttggtgttccactctattttcggcttcgggcaaatccgatcaaaactattctcgacaatcaaaagcgcctggacagtaaagggaatattaaacgctgtcgggttccgttaataaaagaagcagaacaaatcgcgtggttgcaacgtaaattgggcaatgcggcgcgcgttgaagatgtgcatcccatatcggaacggccacagtatttttctggtgatggtaaaagtggaaagatccaaacggtttgctttgaaggtgtgctcaccatcaacgacgcgccagcgttaatagatcttgtacagcaaggtattgggccagctaaatcgatgggatgtggcttgctatctttggctccactgCTCGAGggaGGCGGAGGCGGAAGCGGCGGAGGAGGAAGCGGCGGAGGCGGAAGCggcgGAATTCagagcGGCGGAGGAGGAAGCGGCGGAGGAGGCAGCGGCGGAGGAGGAAGCggGTCGACGGGCGGAGGCGGAAGCGGCGGAGGAGGCAGCGGCGGAGGCGGAAGCggGGTACCtGGCGGAGGAGGCAGCGGCGGAGGAGGAAGCGGCGGAGGAGGAAGCGGCGGAGGAGGCAGCggGGATCCACCGGTCGCCACCATGGTGAGCAAGGGCGAGGAGCTGTTCACCGGGGTGGTGCCCATCCTGGTCGAGCTGGACGGCGACGTAAACGGCCACAAGTTCAGCGTGTCCGGCGAGGGCGAGGGCGATGCCACCTACGGCAAGCTGACCCTGAAGTTCATCTGCACCACCGGCAAGCTGCCCGTGCCCTGGCCCACCCTCGTGACCACCCTGACCTACGGCGTGCAGTGCTTCAGCCGCTACCCCGACCACATGAAGCAGCACGACTTCTTCAAGTCCGCCATGCCCGAAGGCTACGTCCAGGAGCGCACCATCTTCTTCAAGGACGACGGCAACTACAAGACCCGCGCCGAGGTGAAGTTCGAGGGCGACACCCTGGTGAACCGCATCGAGCTGAAGGGCATCGACTTCAAGGAGGACGGCAACATCCTGGGGCACAAGCTGGAGTACAACTACAACAGCCACAACGTCTATATCATGGCCGACAAGCAGAAGAACGGCATCAAGGTGAACTTCAAGATCCGCCACAACATCGAGGACGGCAGCGTGCAGCTCGCCGACCACTACCAGCAGAACACCCCCATCGGCGACGGCCCCGTGCTGCTGCCCGACAACCACTACCTGAGCACCCAGTCCGCCCTGAGCAAAGACCCCAACGAGAAGCGCGATCACATGGTCCTGCTGGAGTTCGTGACCGCCGCCGGGATCACTCTCGGCATGGACGAGCTGTACAAGTAA

Their deduced amino acid sequences are:

MVYLSKVIIARAWSRDLYQLHQGLWHLFPNRPDAARDFLFHVEKRNTPEGCHVLLQSAQMPVSTAVATVIKTKQVEFQLQVGVPLYFRLRANPIKTILDNQKRLDSKGNIKRCRVPLIKEAEQIAWLQRKLGNAARVEDVHPISERPQYFSGDGKSGKIQTVCFEGVLTINDAPALIDLVQQGIGPAKSMGCGLLSLAPLLEGGGGGSGGGGSGGGGSGGIQSGGGGSGGGGSGGGGSGSTGGGGSGGGGSGGGGSGVPGGGGSGGGGSGGGGSGGGGSGDPPVATMVSKGEELFTGVVPILVELDGDVNGHKFSVSGEGEGDATYGKLTLKFICTTGKLPVPWPTLVTTLTYGVQCFSRYPDHMKQHDFFKSAMPEGYVQERTIFFKDDGNYKTRAEVKFEGDTLVNRIELKGIDFKEDGNILGHKLEYNYNSHNVYIMADKQKNGIKVNFKIRHNIEDGSVQLADHYQQNTPIGDGPVLLPDNHYLSTQSALSKDPNEKRDHMVLLEFVTAAGITLGMDELYK*

**Supplementary Figure S2**

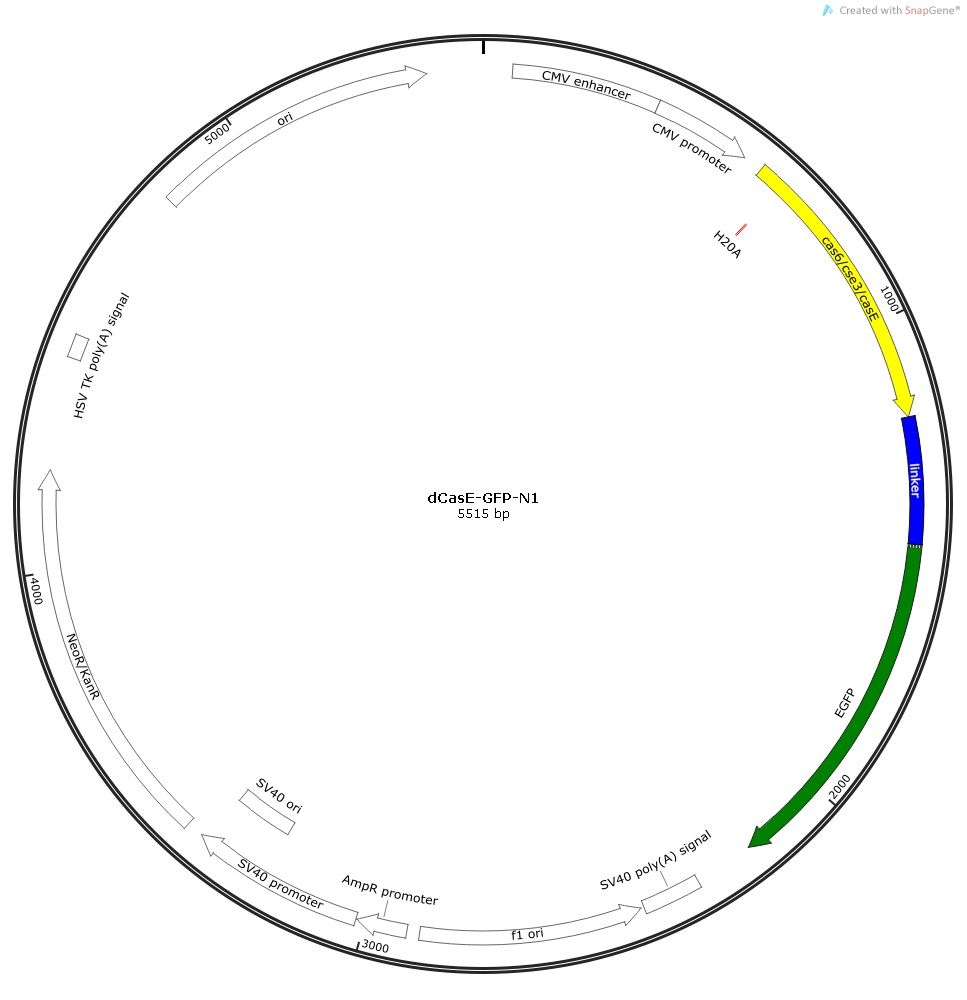

**Supplementary Figure S2.** d*Cas*E-GFP-N1 construct.

The coding sequences of d*Cas*E, linker, and GFP are shown in yellow, blue, and green, respectively. The mutation site His20Ala is marked in red.

ATGGTGtatctcagtaaagtcatcattgccagggcctggagcagggatctttaccaacttgcccagggattatggcatttatttccaaacagaccggatgctgctcgtgattttctttttcatgttgagaagcgaaacacaccagaaggctgtcatgttttattgcagtcagcgcaaatgcctgtttcaactgccgttgcgacagtcattaaaactaaacaggttgaatttcaacttcaggttggtgttccactctattttcggcttcgggcaaatccgatcaaaactattctcgacaatcaaaagcgcctggacagtaaagggaatattaaacgctgtcgggttccgttaataaaagaagcagaacaaatcgcgtggttgcaacgtaaattgggcaatgcggcgcgcgttgaagatgtgcatcccatatcggaacggccacagtatttttctggtgatggtaaaagtggaaagatccaaacggtttgctttgaaggtgtgctcaccatcaacgacgcgccagcgttaatagatcttgtacagcaaggtattgggccagctaaatcgatgggatgtggcttgctatctttggctccactgCTCGAGggaGGCGGAGGCGGAAGCGGCGGAGGAGGAAGCGGCGGAGGCGGAAGCggcgGAATTCagagcGGCGGAGGAGGAAGCGGCGGAGGAGGCAGCGGCGGAGGAGGAAGCggGTCGACGGGCGGAGGCGGAAGCGGCGGAGGAGGCAGCGGCGGAGGCGGAAGCggGGTACCtGGCGGAGGAGGCAGCGGCGGAGGAGGAAGCGGCGGAGGAGGAAGCGGCGGAGGAGGCAGCggGGATCCACCGGTCGCCACCATGGTGAGCAAGGGCGAGGAGCTGTTCACCGGGGTGGTGCCCATCCTGGTCGAGCTGGACGGCGACGTAAACGGCCACAAGTTCAGCGTGTCCGGCGAGGGCGAGGGCGATGCCACCTACGGCAAGCTGACCCTGAAGTTCATCTGCACCACCGGCAAGCTGCCCGTGCCCTGGCCCACCCTCGTGACCACCCTGACCTACGGCGTGCAGTGCTTCAGCCGCTACCCCGACCACATGAAGCAGCACGACTTCTTCAAGTCCGCCATGCCCGAAGGCTACGTCCAGGAGCGCACCATCTTCTTCAAGGACGACGGCAACTACAAGACCCGCGCCGAGGTGAAGTTCGAGGGCGACACCCTGGTGAACCGCATCGAGCTGAAGGGCATCGACTTCAAGGAGGACGGCAACATCCTGGGGCACAAGCTGGAGTACAACTACAACAGCCACAACGTCTATATCATGGCCGACAAGCAGAAGAACGGCATCAAGGTGAACTTCAAGATCCGCCACAACATCGAGGACGGCAGCGTGCAGCTCGCCGACCACTACCAGCAGAACACCCCCATCGGCGACGGCCCCGTGCTGCTGCCCGACAACCACTACCTGAGCACCCAGTCCGCCCTGAGCAAAGACCCCAACGAGAAGCGCGATCACATGGTCCTGCTGGAGTTCGTGACCGCCGCCGGGATCACTCTCGGCATGGACGAGCTGTACAAGTAA

Their deduced amino acid sequences are:

MVYLSKVIIARAWSRDLYQLAQGLWHLFPNRPDAARDFLFHVEKRNTPEGCHVLLQSAQMPVSTAVATVIKTKQVEFQLQVGVPLYFRLRANPIKTILDNQKRLDSKGNIKRCRVPLIKEAEQIAWLQRKLGNAARVEDVHPISERPQYFSGDGKSGKIQTVCFEGVLTINDAPALIDLVQQGIGPAKSMGCGLLSLAPLLEGGGGGSGGGGSGGGGSGGIQSGGGGSGGGGSGGGGSGSTGGGGSGGGGSGGGGSGVPGGGGSGGGGSGGGGSGGGGSGDPPVATMVSKGEELFTGVVPILVELDGDVNGHKFSVSGEGEGDATYGKLTLKFICTTGKLPVPWPTLVTTLTYGVQCFSRYPDHMKQHDFFKSAMPEGYVQERTIFFKDDGNYKTRAEVKFEGDTLVNRIELKGIDFKEDGNILGHKLEYNYNSHNVYIMADKQKNGIKVNFKIRHNIEDGSVQLADHYQQNTPIGDGPVLLPDNHYLSTQSALSKDPNEKRDHMVLLEFVTAAGITLGMDELYK*

**Supplementary Figure S3**

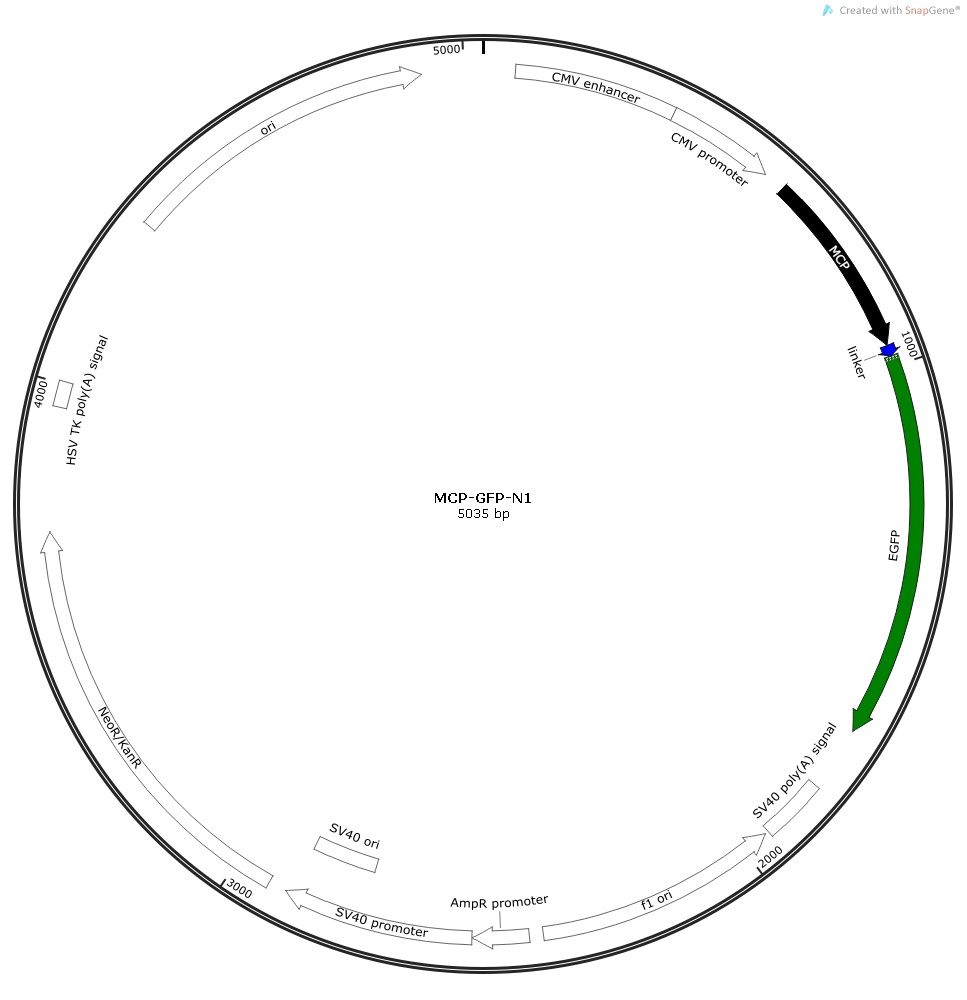

**Supplementary Figure S3.** MCP-GFP-N1 construct.

The coding sequences of MCP, linker, and GFP are shown in black, blue, and green, respectively.

AtggtggcttctaactttactcagttcgttctcgtcgacaatggcggaactggcgacgtgactgtcgccccaagcaacttcgctaacgggatcgctgaatggatcagctctaactcgcgttcacaggcttacaaagtaacctgtagcgttcgtcagagctctgcgcagaatcgcaaatacaccatcaaagtcgaggtgcctaaaggcgcctggcgttcgtacttaaatatggaactaaccattccaattttcgccacgaattccgactgcgagcttattgttaaggcaatgcaaggtctcctaaaagatggaaacccgattccctcagcaatcgcagcaaactccggcatctacgccCTCGAGgcACCGGTCGCCACCATGGTGAGCAAGGGCGAGGAGCTGTTCACCGGGGTGGTGCCCATCCTGGTCGAGCTGGACGGCGACGTAAACGGCCACAAGTTCAGCGTGTCCGGCGAGGGCGAGGGCGATGCCACCTACGGCAAGCTGACCCTGAAGTTCATCTGCACCACCGGCAAGCTGCCCGTGCCCTGGCCCACCCTCGTGACCACCCTGACCTACGGCGTGCAGTGCTTCAGCCGCTACCCCGACCACATGAAGCAGCACGACTTCTTCAAGTCCGCCATGCCCGAAGGCTACGTCCAGGAGCGCACCATCTTCTTCAAGGACGACGGCAACTACAAGACCCGCGCCGAGGTGAAGTTCGAGGGCGACACCCTGGTGAACCGCATCGAGCTGAAGGGCATCGACTTCAAGGAGGACGGCAACATCCTGGGGCACAAGCTGGAGTACAACTACAACAGCCACAACGTCTATATCATGGCCGACAAGCAGAAGAACGGCATCAAGGTGAACTTCAAGATCCGCCACAACATCGAGGACGGCAGCGTGCAGCTCGCCGACCACTACCAGCAGAACACCCCCATCGGCGACGGCCCCGTGCTGCTGCCCGACAACCACTACCTGAGCACCCAGTCCGCCCTGAGCAAAGACCCCAACGAGAAGCGCGATCACATGGTCCTGCTGGAGTTCGTGACCGCCGCCGGGATCACTCTCGGCATGGACGAGCTGTACAAGTAA

Their deduced amino acid sequences are:

MVASNFTQFVLVDNGGTGDVTVAPSNFANGIAEWISSNSRSQAYKVTCSVRQSSAQNRKYTIKVEVPKGAWRSYLNMELTIPIFATNSDCELIVKAMQGLLKDGNPIPSAIAANSGIYALEAPVATMVSKGEELFTGVVPILVELDGDVNGHKFSVSGEGEGDATYGKLTLKFICTTGKLPVPWPTLVTTLTYGVQCFSRYPDHMKQHDFFKSAMPEGYVQERTIFFKDDGNYKTRAEVKFEGDTLVNRIELKGIDFKEDGNILGHKLEYNYNSHNVYIMADKQKNGIKVNFKIRHNIEDGSVQLADHYQQNTPIGDGPVLLPDNHYLSTQSALSKDPNEKRDHMVLLEFVTAAGITLGMDELYK*

**Supplementary Figure S4**

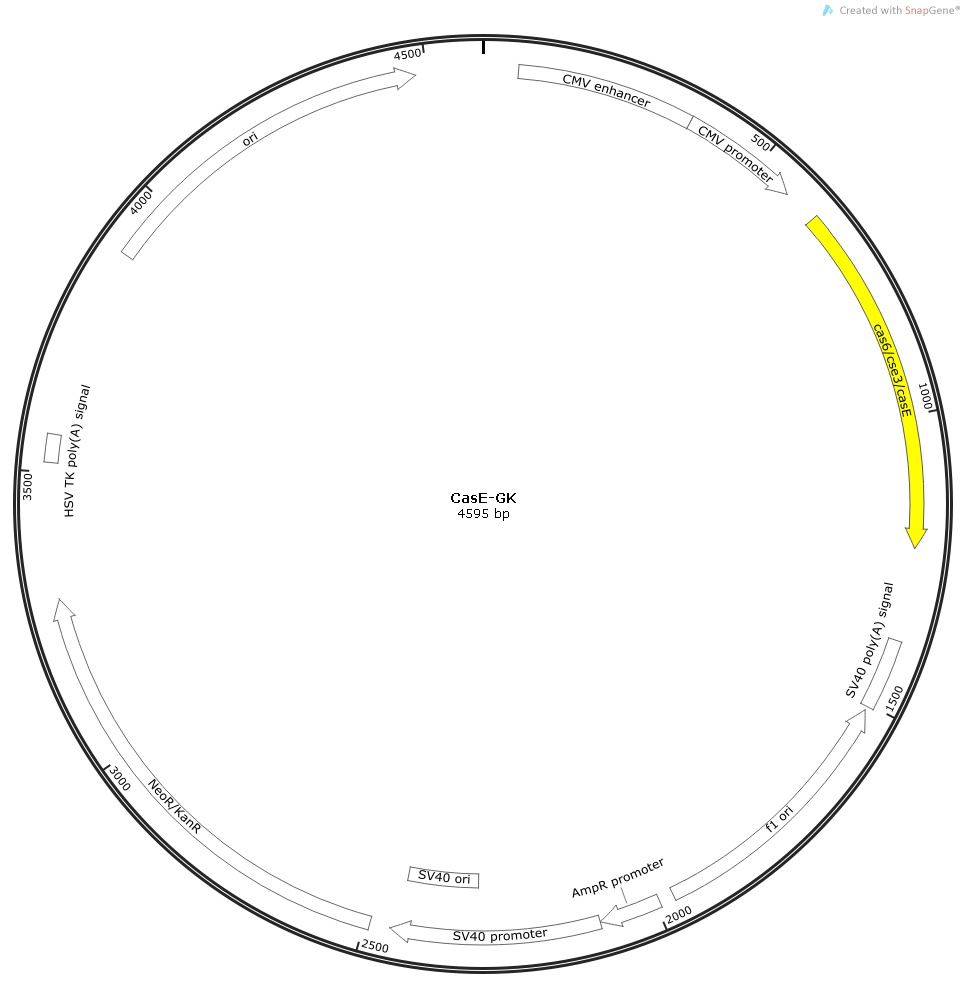

**Supplementary Figure S4.** *Cas*E-GK construct.

The coding sequence of *Cas*E is marked in yellow.

GGTTTAGTGAACCGTCAGATCCGCTAGCGCTACCGGACTCAGATCTCGAGGCCACCATGtatctcagtaaagtcatcattgccagggcctggagcagggatctttaccaacttcaccagggattatggcatttatttccaaacagaccggatgctgctcgtgattttctttttcatgttgagaagcgaaacacaccagaaggctgtcatgttttattgcagtcagcgcaaatgcctgtttcaactgccgttgcgacagtcattaaaactaaacaggttgaatttcaacttcaggttggtgttccactctattttcggcttcgggcaaatccgatcaaaactattctcgacaatcaaaagcgcctggacagtaaagggaatattaaacgctgtcgggttccgttaataaaagaagcagaacaaatcgcgtggttgcaacgtaaattgggcaatgcggcgcgcgttgaagatgtgcatcccatatcggaacggccacagtatttttctggtgatggtaaaagtggaaagatccaaacggtttgctttgaaggtgtgctcaccatcaacgacgcgccagcgttaatagatcttgtacagcaaggtattgggccagctaaatcgatgggatgtggcttgctatctttggctccactgtgaGGTACCGCGGGCCCGGGATCCATATGTAAGTAAGTAAGCGGCCGCGACTCTAGATCATAATCAGCCATACCACATTTGTAGAGGTTTTACTTGCTTTAAAAAACCTCCCACACCTCCCCCTGAACCTGAAACATAAAATGAATGCAATTGTTGTTGTT

**Supplementary Figure S5**

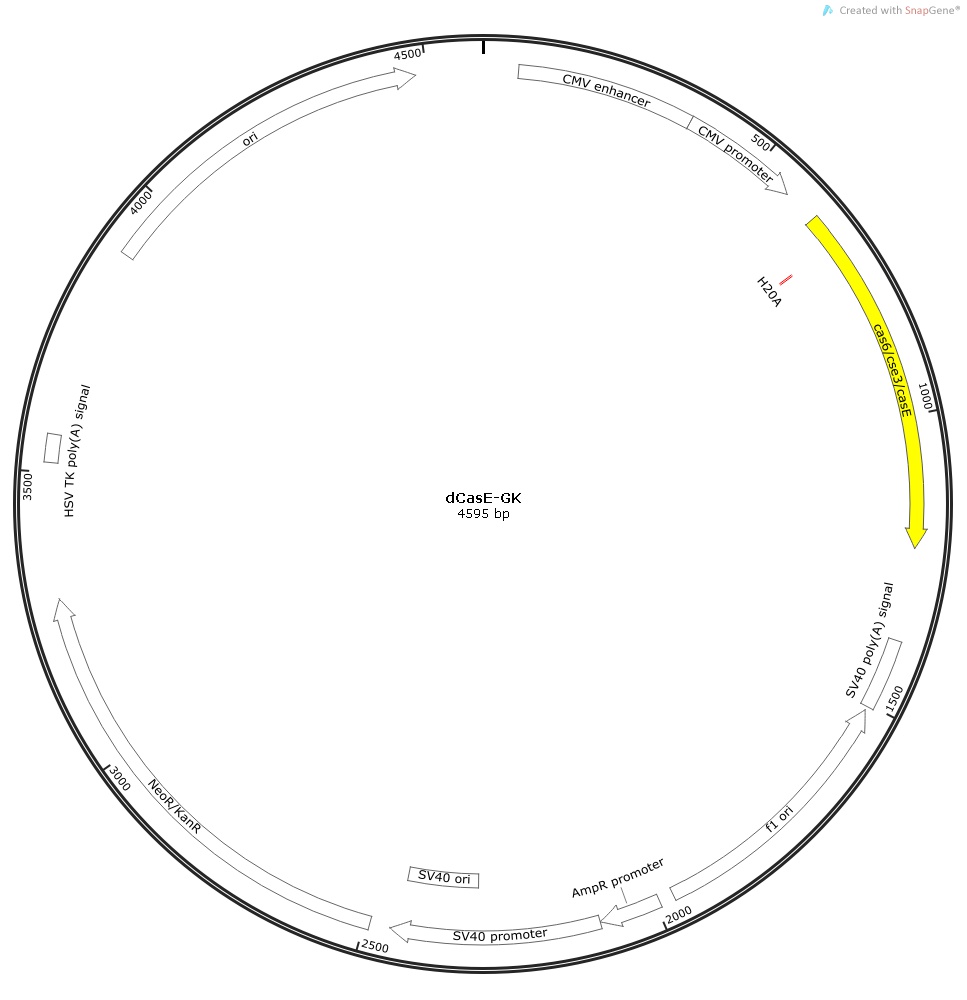

**Supplementary Figure S5.** d*Cas*E-GK construct.

The coding sequence of d*Cas*E is marked in yellow. The mutation site His20Ala is marked in red.

GGTTTAGTGAACCGTCAGATCCGCTAGCGCTACCGGACTCAGATCTCGAGGCCACCATGtatctcagtaaagtcatcattgccagggcctggagcagggatctttaccaacttgcccagggattatggcatttatttccaaacagaccggatgctgctcgtgattttctttttcatgttgagaagcgaaacacaccagaaggctgtcatgttttattgcagtcagcgcaaatgcctgtttcaactgccgttgcgacagtcattaaaactaaacaggttgaatttcaacttcaggttggtgttccactctattttcggcttcgggcaaatccgatcaaaactattctcgacaatcaaaagcgcctggacagtaaagggaatattaaacgctgtcgggttccgttaataaaagaagcagaacaaatcgcgtggttgcaacgtaaattgggcaatgcggcgcgcgttgaagatgtgcatcccatatcggaacggccacagtatttttctggtgatggtaaaagtggaaagatccaaacggtttgctttgaaggtgtgctcaccatcaacgacgcgccagcgttaatagatcttgtacagcaaggtattgggccagctaaatcgatgggatgtggcttgctatctttggctccactgtgaGGTACCGCGGGCCCGGGATCCATATGTAAGTAAGTAAGCGGCCGCGACTCTAGATCATAATCAGCCATACCACATTTGTAGAGGTTTTACTTGCTTTAAAAAACCTCCCACACCTCCCCCTGAACCTGAAACATAAAATGAATGCAATTGTTGTTGTT

**Supplementary Figure S6**

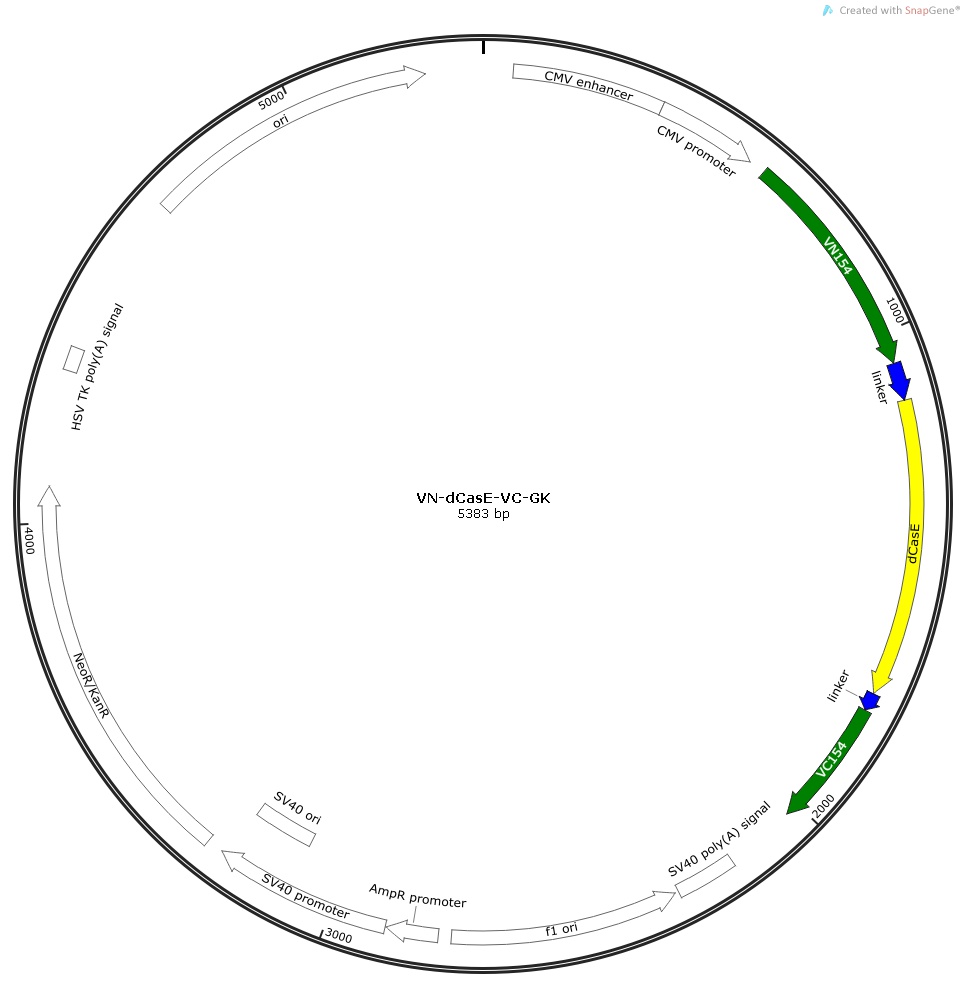

**Supplementary Figure S6.** VN-d*Cas*E-VC-GK construct.

The nucleotide sequences of the Venus N-terminus (1‒154 aa) and C-terminus (155‒240 aa) , linker, d*Cas*E, and the mutation site His20Ala are marked in green, blue, yellow, and red, respectively.

ATGGTGAGCAAGGGCGAGGAGCTGTTCACCGGGGTGGTGCCCATCCTGGTCGAGCTGGACGGCGACGTAAACGGCCACAAGTTCAGCGTGTCCGGCGAGGGCGAGGGCGATGCCACCTACGGCAAGCTGACCCTGAAGCTGATCTGCACCACCGGCAAGCTGCCCGTGCCCTGGCCCACCCTCGTGACCACCCTGGGCTACGGCCTGCAGTGCTTCGCCCGCTACCCCGACCACATGAAGCAGCACGACTTCTTCAAGTCCGCCATGCCCGAAGGCTACGTCCAGGAGCGCACCATCTTCTTCAAGGACGACGGCAACTACAAGACCCGCGCCGAGGTGAAGTTCGAGGGCGACACCCTGGTGAACCGCATCGAGCTGAAGGGCATCGACTTCAAGGAGGACGGCAACATCCTGGGGCACAAGCTGGAGTACAACTACAACAGCCACAACGTCTATATCACCCTCGAGAGACCTGCTTGTAAAATTCCAAACGACCTGAAGCAGAAAGTGATGAACCACAAGCTTGCCACCATGGTGTATCTCAGTAAAGTCATCATTGCCAGGGCCTGGAGCAGGGATCTTTACCAACTTgcccagggattatggcatttatttccaaacagaccggatgctgctcgtgattttctttttcatgttgagaagcgaaacacaccagaaggctgtcatgttttattgcagtcagcgcaaatgcctgtttcaactgccgttgcgacagtcattaaaactaaacaggttgaatttcaacttcaggttggtgttccactctattttcggcttcgggcaaatccgatcaaaactattctcgacaatcaaaagcgcctggacagtaaagggaatattaaacgctgtcgggttccgttaataaaagaagcagaacaaatcgcgtggttgcaacgtaaattgggcaatgcggcgcgcgttgaagatgtgcatcccatatcggaacggccacagtatttttctggtgatggtaaaagtggaaagatccaaacggtttgctttgaaggtgtgctcaccatcaacgacgcgccagcgttaatagatcttgtacagcaaggtattgggccagctaaatcgatgggatgtggcttgctatctttggctccactGCTGCAGTCGACGACCTGCACAGCTGGCGCTGAATTCGCCGACAAGCAGAAGAACGGCATCAAGGCCAACTTCAAGATCCGCCACAACATCGAGGACGGCGGCGTGCAGCTCGCCGACCACTACCAGCAGAACACCCCCATCGGCGACGGCCCCGTGCTGCTGCCCGACAACCACTACCTGAGCTACCAGTCCGCCCTGAGCAAAGACCCCAACGAGAAGCGCGATCACATGGTCCTGCTGGAGTTCGTGACCGCCGCCGGGATCACTCTCGGCATGGACGAGCTGTACAAGTAA

Their deduced amino acid sequences are:

MVSKGEELFTGVVPILVELDGDVNGHKFSVSGEGEGDATYGKLTLKLICTTGKLPVPWPTLVTTLGYGLQCFARYPDHMKQHDFFKSAMPEGYVQERTIFFKDDGNYKTRAEVKFEGDTLVNRIELKGIDFKEDGNILGHKLEYNYNSHNVYITLERPACKIPNDLKQKVMNHKLATMVYLSKVIIARAWSRDLYQLAQGLWHLFPNRPDAARDFLFHVEKRNTPEGCHVLLQSAQMPVSTAVATVIKTKQVEFQLQVGVPLYFRLRANPIKTILDNQKRLDSKGNIKRCRVPLIKEAEQIAWLQRKLGNAARVEDVHPISERPQYFSGDGKSGKIQTVCFEGVLTINDAPALIDLVQQGIGPAKSMGCGLLSLAPLLQSTTCTAGAEFADKQKNGIKANFKIRHNIEDGGVQLADHYQQNTPIGDGPVLLPDNHYLSYQSALSKDPNEKRDHMVLLEFVTAAGITLGMDELYK*

**Supplementary Figure S7**

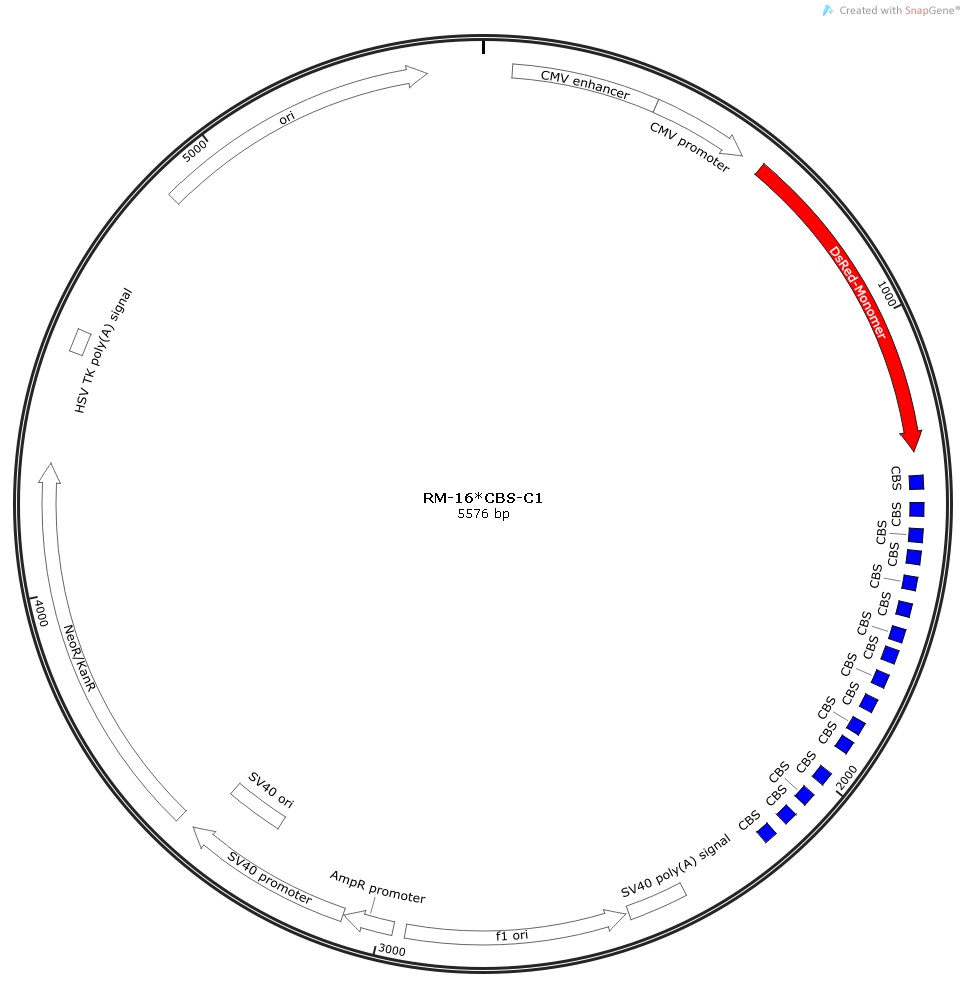

**Supplementary Figure S7.** RM-16×CBS-C1 construct.

The nucleotide sequences of RFP monomer and CBS are marked in red and blue, respectively.

GGTTTAGTGAACCGTCAGATCCGCTAGCGCTACCGGTCGCCACCatgGACAACACCGAGGACGTCATCAAGGAGTTCATGCAGTTCAAGGTGCGCATGGAGGGCTCCGTGAACGGCCACTACTTCGAGATCGAGGGCGAGGGCGAGGGCAAGCCCTACGAGGGCACCCAGACCGCCAAGCTGCAGGTGACCAAGGGCGGCCCCCTGCCCTTCGCCTGGGACATCCTGTCCCCCCAGTTCCAGTACGGCTCCAAGGCCTACGTGAAGCACCCCGCCGACATCCCCGACTACATGAAGCTGTCCTTCCCCGAGGGCTTCACCTGGGAGCGCTCCATGAACTTCGAGGACGGCGGCGTGGTGGAGGTGCAGCAGGACTCCTCCCTGCAGGACGGCACCTTCATCTACAAGGTGAAGTTCAAGGGCGTGAACTTCCCCGCCGACGGCCCCGTAATGCAGAAGAAGACTGCCGGCTGGGAGCCCTCCACCGAGAAGCTGTACCCCCAGGACGGCGTGCTGAAGGGCGAGATCTCCCACGCCCTGAAGCTGAAGGACGGCGGCCACTACACCTGcGACTTCAAGACCGTGTACAAGGCCAAGAAGCCCGTGCAGCTGCCCGGCAACCACTACGTGGACTCCAAGCTGGACATCACCAACCACAACGAGGACTACACCGTGGTGGAGCAGTACGAGCACGCCGAGGCCCGCCACTCCGGCTCCCAGTCCGGACTCAGATCTCGACTTTGAGCGCTACCGGACTCAGATCTCGACCGAGTTCCCCGCGCCAGCGGGGATAAACCGCGCCAGCTTCGAATTCTGCAGTCGAAGAGTTCCCCGCGCCAGCGGGGATAAACCCGATCCACCGGTATTTATCTCGACCGAGTTCCCCGCGCCAGCGGGGATAAACCGCGCCAGCTTCGAATTCGAGTTCCCCGCGCCAGCGGGGATAAACCCGATCCACCGGTATTTATCTCGACCGAGTTCCCCGCGCCAGCGGGGATAAACCGCGCCAGCTTCGAATTCTGCAGTCGAAGAGTTCCCCGCGCCAGCGGGGATAAACCCGATCCACCGGTATTTATCTCGACCGAGTTCCCCGCGCCAGCGGGGATAAACCGCGCCAGCTTCGAATTCGAGTTCCCCGCGCCAGCGGGGATAAACCCGATCCACCGGTATTTATCTCGACCGAGTTCCCCGCGCCAGCGGGGATAAACCGCGCCAGCTTCGAATTCTGCAGTCGAAGAGTTCCCCGCGCCAGCGGGGATAAACCCGATCCACCGGTATTTATCTCGACCGAGTTCCCCGCGCCAGCGGGGATAAACCGCGCCAGCTTCGAATTCGAGTTCCCCGCGCCAGCGGGGATAAACCCGATCCACCGGTATTTATCTCGAGGGAAGCTTCGAATTCTGCAGTCGACCGAGTTCCCCGCGCCAGCGGGGATAAACCGCGCCAGCTTCGAATTCTGCAGTCGAAGAGTTCCCCGCGCCAGCGGGGATAAACCCGATCCACCGGTATTTATCTCGACCGAGTTCCCCGCGCCAGCGGGGATAAACCGCGCCAGCTTCGAATTCTGCAGTCGAAGAGTTCCCCGCGCCAGCGGGGATAAACCCGATCCACCGGTATTTATCTCGAGGGAAGCTTCGAATTCTGCAGTCGACGGTACCGCGGGCCCgGGATCCACCGGATCTAGATAACTGATCATAATCAGCCATACCACATTTGTAGAGGTTTTACTTGCTTTAAAAAACCTCCCACACCTCCCCCTGAACCTGAAACATAAAATGAATGCAATTGTTGTTGTT

**Supplementary Figure S8**

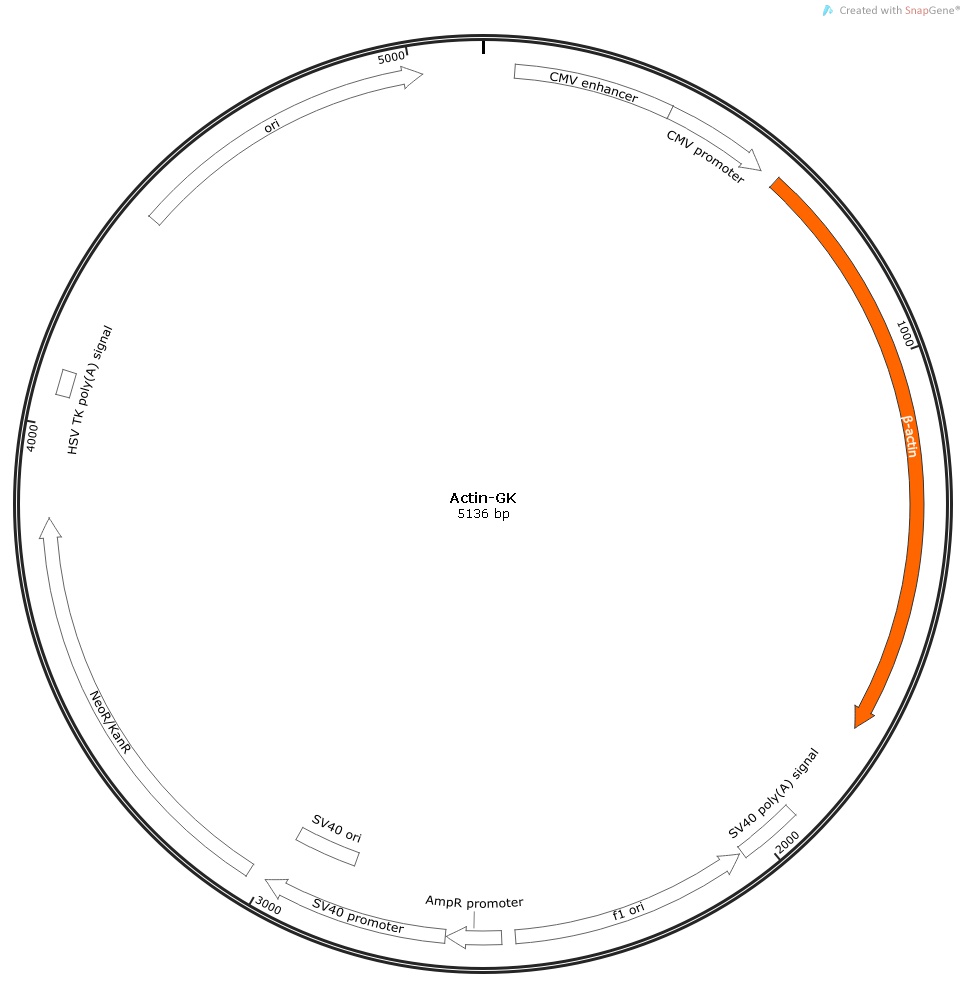

**Supplementary Figure S8.** Actin-GK construct.

The β-actin ORF is marked in orange.

GGTTTAGTGAACCGTCAGATCCGCTAGCGCCACCatggatgatgatatcgccgcgctcgtcgtcgacaacggctccggcatgtgcaaggccggcttcgcgggcgacgatgccccccgggccgtcttcccctccatcgtggggcgccccaggcaccagggcgtgatggtgggcatgggtcagaaggattcctatgtgggcgacgaggcccagagcaagagaggcatcctcaccctgaagtaccccatcgagcacggcatcgtcaccaactgggacgacatggagaaaatctggcaccacaccttctacaatgagctgcgtgtggctcccgaggagcaccccgtgctgctgaccgaggcccccctgaaccccaaggccaaccgcgagaagatgacccagatcatgtttgagaccttcaacaccccagccatgtacgttgctatccaggctgtgctatccctgtacgcctctggccgtaccactggcatcgtgatggactccggtgacggggtcacccacactgtgcccatctacgaggggtatgccctcccccatgccatcctgcgtctggacctggctggccgggacctgactgactacctcatgaagatcctcaccgagcgcggctacagcttcaccaccacggccgagcgggaaatcgtgcgtgacattaaggagaagctgtgctacgtcgccctggacttcgagcaagagatggccacggctgcttccagctcctccctggagaagagctacgagctgcctgacggccaggtcatcaccattggcaatgagcggttccgctgccctgaggcactcttccagccttccttcctgggcatggagtcctgtggcatccacgaaactaccttcaactccatcatgaagtgtgacgtggacatccgcaaagacctgtacgccaacacagtgctgtctggcggcaccaccatgtaccctggcattgccgacaggatgcagaaggagatcactgccctggcacccagcacaatgaagatcaagatcattgctcctcctgagcgcaagtactccgtgtggatcggcggctccatcctggcctcgctgtccaccttccagcagatgtggatcagcaagcaggagtatgacgagtccggcccctccatcgtccaccgcaaatgcttctgAGATCTCGAGCTCAAGCTTCGAATTCTGCAGTCGACGGTACCGCGGGCCCGGGATCCATATGTAAGTAAGTAAGCGGCCGCGACTCTAGATCATAATCAGCCATACCACATTTGTAGAGGTTTTACTTGCTTTAAAAAACCTCCCACACCTCCCCCTGAACCTGAAACATAAAATGAATGCAATTGTTGTTGTT

**Supplementary Figure S9**

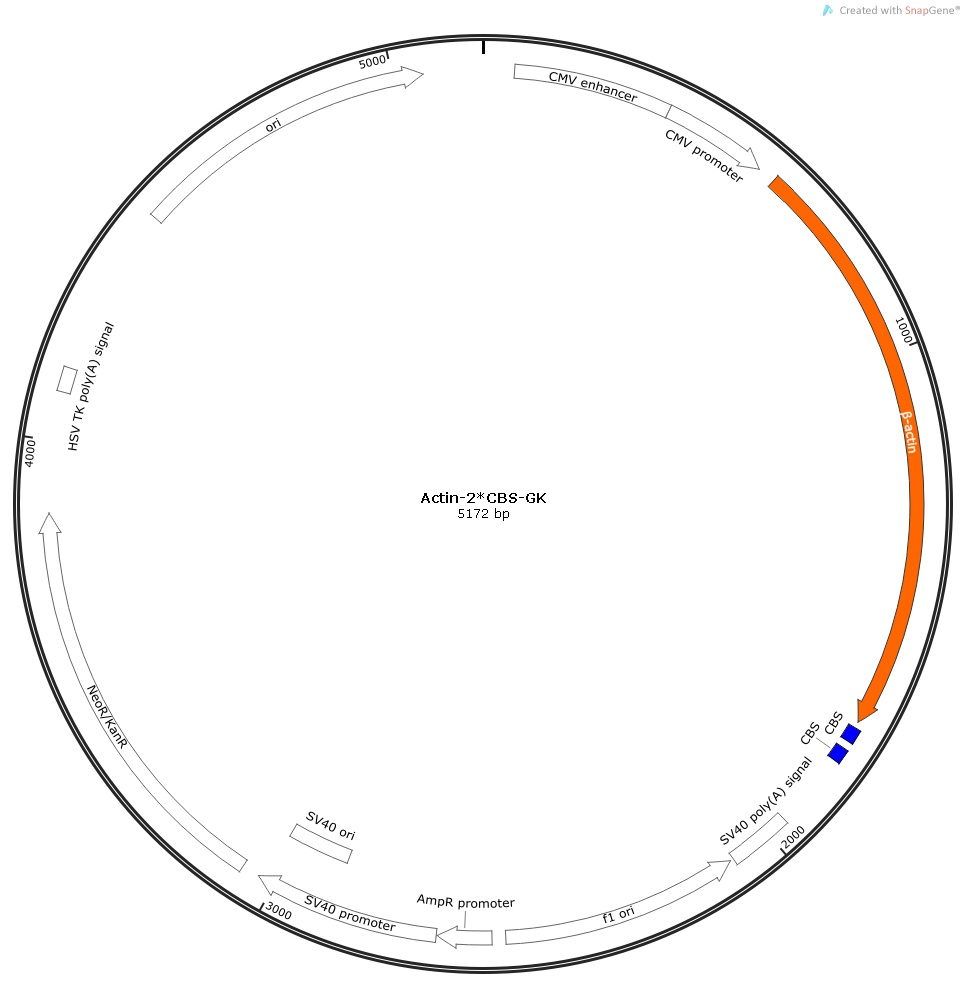

**Supplementary Figure S9.** Actin-2×CBS-GK construct.

The nucleotide sequences of β-actin and CBS are marked in orange and blue, respectively.

GGTTTAGTGAACCGTCAGATCCGCTAGCGCCACCatggatgatgatatcgccgcgctcgtcgtcgacaacggctccggcatgtgcaaggccggcttcgcgggcgacgatgccccccgggccgtcttcccctccatcgtggggcgccccaggcaccagggcgtgatggtgggcatgggtcagaaggattcctatgtgggcgacgaggcccagagcaagagaggcatcctcaccctgaagtaccccatcgagcacggcatcgtcaccaactgggacgacatggagaaaatctggcaccacaccttctacaatgagctgcgtgtggctcccgaggagcaccccgtgctgctgaccgaggcccccctgaaccccaaggccaaccgcgagaagatgacccagatcatgtttgagaccttcaacaccccagccatgtacgttgctatccaggctgtgctatccctgtacgcctctggccgtaccactggcatcgtgatggactccggtgacggggtcacccacactgtgcccatctacgaggggtatgccctcccccatgccatcctgcgtctggacctggctggccgggacctgactgactacctcatgaagatcctcaccgagcgcggctacagcttcaccaccacggccgagcgggaaatcgtgcgtgacattaaggagaagctgtgctacgtcgccctggacttcgagcaagagatggccacggctgcttccagctcctccctggagaagagctacgagctgcctgacggccaggtcatcaccattggcaatgagcggttccgctgccctgaggcactcttccagccttccttcctgggcatggagtcctgtggcatccacgaaactaccttcaactccatcatgaagtgtgacgtggacatccgcaaagacctgtacgccaacacagtgctgtctggcggcaccaccatgtaccctggcattgccgacaggatgcagaaggagatcactgccctggcacccagcacaatgaagatcaagatcattgctcctcctgagcgcaagtactccgtgtggatcggcggctccatcctggcctcgctgtccaccttccagcagatgtggatcagcaagcaggagtatgacgagtccggcccctccatcgtccaccgcaaatgcttctgaGATCTGACCGAGTTCCCCGCGCCAGCGGGGATAAACCGCGCCAGCTTCGCGACGAGTTCCCCGCGCCAGCGGGGATAAACCGCGCCGGATCCATATGTAAGTAAGTAAGCGGCCGCGACTCTAGATCATAATCAGCCATACCACATTTGTAGAGGTTTTACTTGCTTTAAAAAACCTCCCACACCTCCCCCTGAACCTGAAACATAAAATGAATGCAATTGTTGTTGTT

**Supplementary Figure S10**

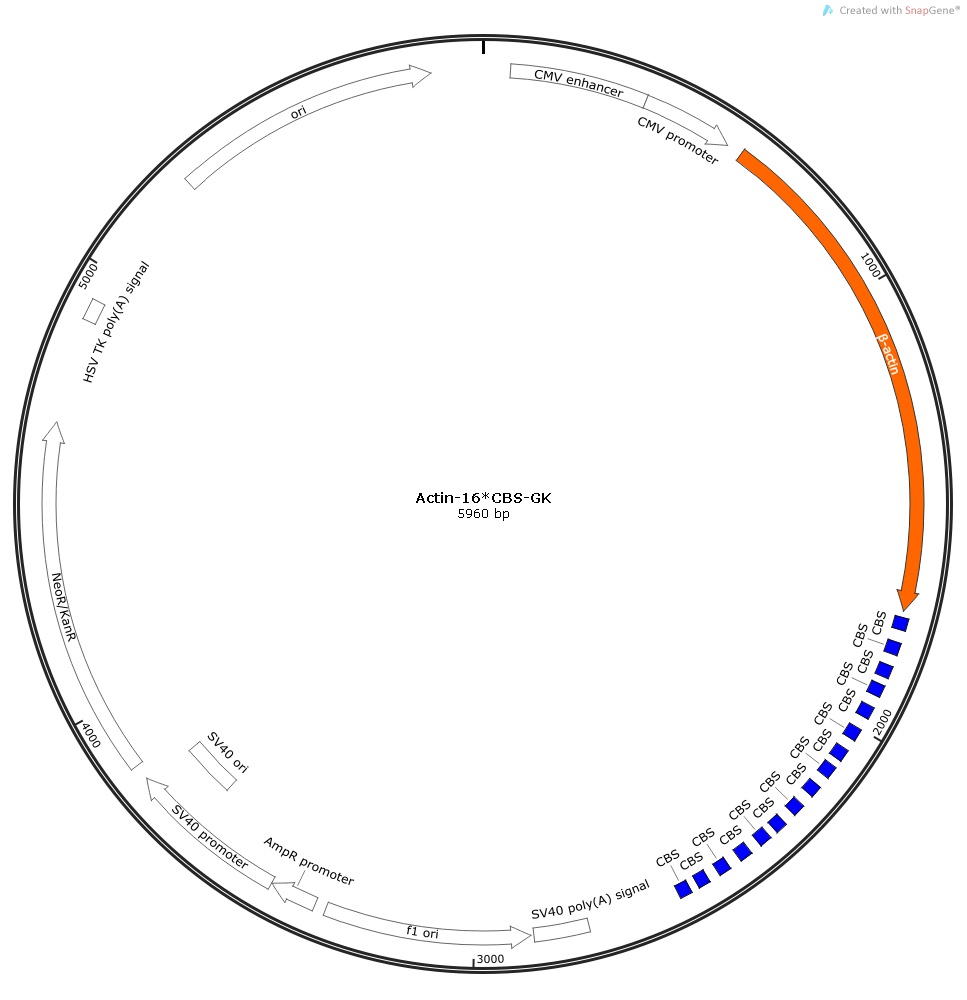

**Supplementary Figure S10.** Actin-16×CBS-GK construct.

The nucleotide sequences of β-actin and CBS are marked in orange and blue, respectively.

GGTTTAGTGAACCGTCAGATCCGCTAGCGCCACCatggatgatgatatcgccgcgctcgtcgtcgacaacggctccggcatgtgcaaggccggcttcgcgggcgacgatgccccccgggccgtcttcccctccatcgtggggcgccccaggcaccagggcgtgatggtgggcatgggtcagaaggattcctatgtgggcgacgaggcccagagcaagagaggcatcctcaccctgaagtaccccatcgagcacggcatcgtcaccaactgggacgacatggagaaaatctggcaccacaccttctacaatgagctgcgtgtggctcccgaggagcaccccgtgctgctgaccgaggcccccctgaaccccaaggccaaccgcgagaagatgacccagatcatgtttgagaccttcaacaccccagccatgtacgttgctatccaggctgtgctatccctgtacgcctctggccgtaccactggcatcgtgatggactccggtgacggggtcacccacactgtgcccatctacgaggggtatgccctcccccatgccatcctgcgtctggacctggctggccgggacctgactgactacctcatgaagatcctcaccgagcgcggctacagcttcaccaccacggccgagcgggaaatcgtgcgtgacattaaggagaagctgtgctacgtcgccctggacttcgagcaagagatggccacggctgcttccagctcctccctggagaagagctacgagctgcctgacggccaggtcatcaccattggcaatgagcggttccgctgccctgaggcactcttccagccttccttcctgggcatggagtcctgtggcatccacgaaactaccttcaactccatcatgaagtgtgacgtggacatccgcaaagacctgtacgccaacacagtgctgtctggcggcaccaccatgtaccctggcattgccgacaggatgcagaaggagatcactgccctggcacccagcacaatgaagatcaagatcattgctcctcctgagcgcaagtactccgtgtggatcggcggctccatcctggcctcgctgtccaccttccagcagatgtggatcagcaagcaggagtatgacgagtccggcccctccatcgtccaccgcaaatgcttctgaGATCTCGACCGAGTTCCCCGCGCCAGCGGGGATAAACCGCGCCAGCTTCGAATTCTGCAGTCGAAGAGTTCCCCGCGCCAGCGGGGATAAACCCGATCCACCGGTATTTATCTCGACCGAGTTCCCCGCGCCAGCGGGGATAAACCGCGCCAGCTTCGAATTCGAGTTCCCCGCGCCAGCGGGGATAAACCCGATCCACCGGTATTTATCTCGACCGAGTTCCCCGCGCCAGCGGGGATAAACCGCGCCAGCTTCGAATTCTGCAGTCGAAGAGTTCCCCGCGCCAGCGGGGATAAACCCGATCCACCGGTATTTATCTCGACCGAGTTCCCCGCGCCAGCGGGGATAAACCGCGCCAGCTTCGAATTCGAGTTCCCCGCGCCAGCGGGGATAAACCCGATCCACCGGTATTTATCTCGACCGAGTTCCCCGCGCCAGCGGGGATAAACCGCGCCAGCTTCGAATTCTGCAGTCGAAGAGTTCCCCGCGCCAGCGGGGATAAACCCGATCCACCGGTATTTATCTCGACCGAGTTCCCCGCGCCAGCGGGGATAAACCGCGCCAGCTTCGAATTCGAGTTCCCCGCGCCAGCGGGGATAAACCCGATCCACCGGTATTTATCTCGACCGAGTTCCCCGCGCCAGCGGGGATAAACCGCGCCAGCTTCGAATTCTGCAGTCGAAGAGTTCCCCGCGCCAGCGGGGATAAACCCGATCCACCGGTATTTATCTCGACCGAGTTCCCCGCGCCAGCGGGGATAAACCGCGCCAGCTTCGAATTCGAGTTCCCCGCGCCAGCGGGGATAAACCCGATCCACCGGTATTTATCTCGAGCTCAAGCTTCGAATTCTGCAGTCGACGGTACCGCGGGCCCGGGATCCATATGTAAGTAAGTAAGCGGCCGCGACTCTAGATCATAATCAGCCATACCACATTTGTAGAGGTTTTACTTGCTTTAAAAAACCTCCCACACCTCCCCCTGAACCTGAAACATAAAATGAATGCAATTGTTGTTGTT

**Supplementary Figure S11**

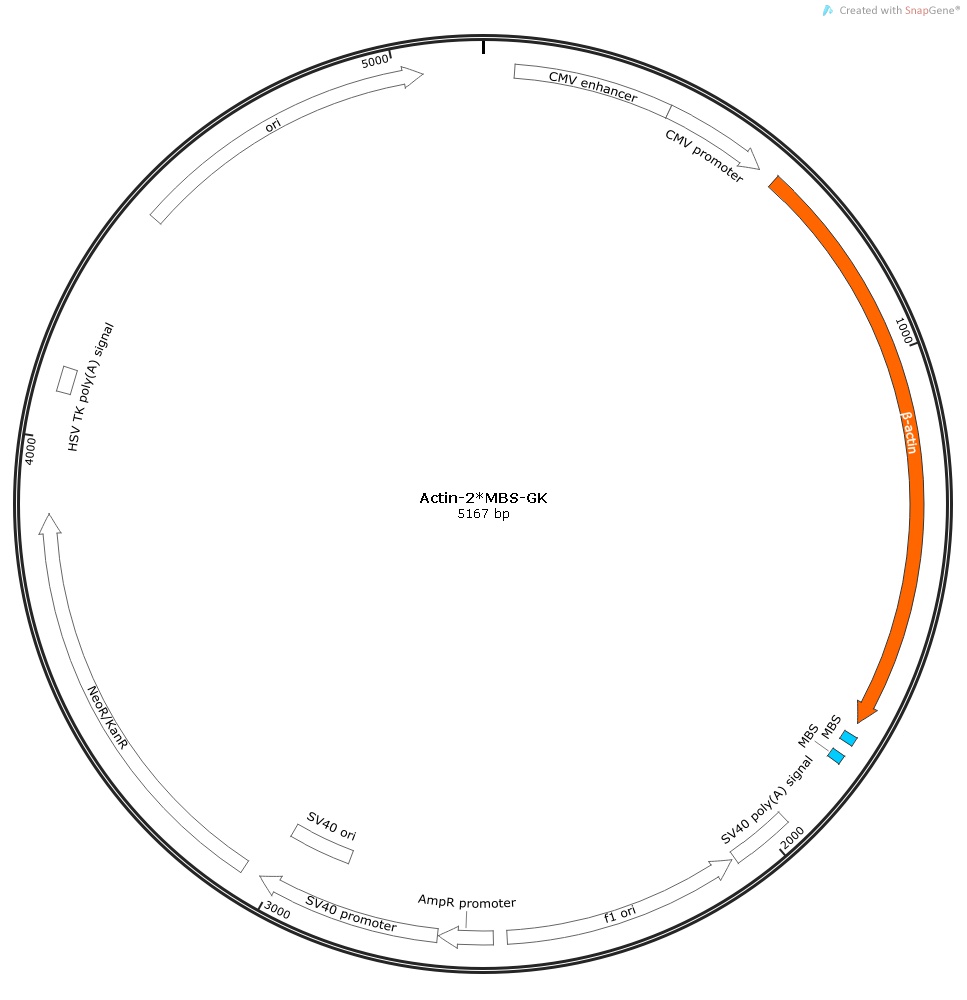

**Supplementary Figure S11.** Actin-2×MBS-GK construct.

The nucleotide sequences of β-actin are marked in orange and blue, respectively

GGTTTAGTGAACCGTCAGATCCGCTAGCGCCACCatggatgatgatatcgccgcgctcgtcgtcgacaacggctccggcatgtgcaaggccggcttcgcgggcgacgatgccccccgggccgtcttcccctccatcgtggggcgccccaggcaccagggcgtgatggtgggcatgggtcagaaggattcctatgtgggcgacgaggcccagagcaagagaggcatcctcaccctgaagtaccccatcgagcacggcatcgtcaccaactgggacgacatggagaaaatctggcaccacaccttctacaatgagctgcgtgtggctcccgaggagcaccccgtgctgctgaccgaggcccccctgaaccccaaggccaaccgcgagaagatgacccagatcatgtttgagaccttcaacaccccagccatgtacgttgctatccaggctgtgctatccctgtacgcctctggccgtaccactggcatcgtgatggactccggtgacggggtcacccacactgtgcccatctacgaggggtatgccctcccccatgccatcctgcgtctggacctggctggccgggacctgactgactacctcatgaagatcctcaccgagcgcggctacagcttcaccaccacggccgagcgggaaatcgtgcgtgacattaaggagaagctgtgctacgtcgccctggacttcgagcaagagatggccacggctgcttccagctcctccctggagaagagctacgagctgcctgacggccaggtcatcaccattggcaatgagcggttccgctgccctgaggcactcttccagccttccttcctgggcatggagtcctgtggcatccacgaaactaccttcaactccatcatgaagtgtgacgtggacatccgcaaagacctgtacgccaacacagtgctgtctggcggcaccaccatgtaccctggcattgccgacaggatgcagaaggagatcactgccctggcacccagcacaatgaagatcaagatcattgctcctcctgagcgcaagtactccgtgtggatcggcggctccatcctggcctcgctgtccaccttccagcagatgtggatcagcaagcaggagtatgacgagtccggcccctccatcgtccaccgcaaatgcttctgaGATCTATATTCGGCCAAGAAAGCACGAGCATCAGCCGTGCCTCCAGGTCGAATCTTCAAATTGCACGAGCATCAGCCGTGCGGATCCATATGTAAGTAAGTAAGCGGCCGCGACTCTAGATCATAATCAGCCATACCACATTTGTAGAGGTTTTACTTGCTTTAAAAAACCTCCCACACCTCCCCCTGAACCTGAAACATAAAATGAATGCAATTGTTGTTGTT

**Supplementary Figure S12**

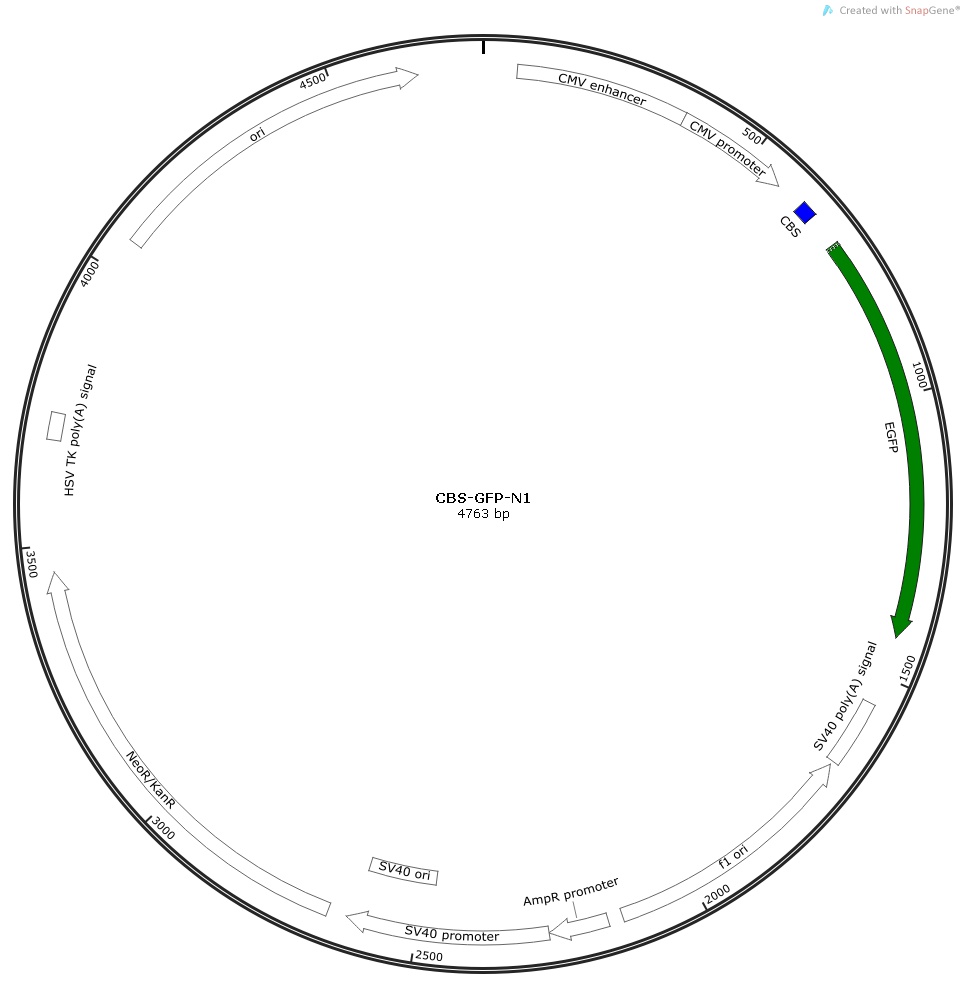

**Supplementary Figure S12.** CBS-GFP-N1 construct.

The nucleotide sequences of CBS and EGFP are marked in blue and green, respectively.

GGTTTAGTGAACCGTCAGATCCGCTAGCGCTACCGGACTCAGATCTCGACCGAGTTCCCCGCGCCAGCGGGGATAAACCGCGCCAGCTTCGAATTCTGCAGTCGACGGTACCGCGGGCCCGGGATCCACCGGTCGCCACCATGGTGAGCAAGGGCGAGGAGCTGTTCACCGGGGTGGTGCCCATCCTGGTCGAGCTGGACGGCGACGTAAACGGCCACAAGTTCAGCGTGTCCGGCGAGGGCGAGGGCGATGCCACCTACGGCAAGCTGACCCTGAAGTTCATCTGCACCACCGGCAAGCTGCCCGTGCCCTGGCCCACCCTCGTGACCACCCTGACCTACGGCGTGCAGTGCTTCAGCCGCTACCCCGACCACATGAAGCAGCACGACTTCTTCAAGTCCGCCATGCCCGAAGGCTACGTCCAGGAGCGCACCATCTTCTTCAAGGACGACGGCAACTACAAGACCCGCGCCGAGGTGAAGTTCGAGGGCGACACCCTGGTGAACCGCATCGAGCTGAAGGGCATCGACTTCAAGGAGGACGGCAACATCCTGGGGCACAAGCTGGAGTACAACTACAACAGCCACAACGTCTATATCATGGCCGACAAGCAGAAGAACGGCATCAAGGTGAACTTCAAGATCCGCCACAACATCGAGGACGGCAGCGTGCAGCTCGCCGACCACTACCAGCAGAACACCCCCATCGGCGACGGCCCCGTGCTGCTGCCCGACAACCACTACCTGAGCACCCAGTCCGCCCTGAGCAAAGACCCCAACGAGAAGCGCGATCACATGGTCCTGCTGGAGTTCGTGACCGCCGCCGGGATCACTCTCGGCATGGACGAGCTGTACAAGTAAAGCGGCCGCGACTCTAGATCATAATCAGCCATACCACATTTGTAGAGGTTTTACTTGCTTTAAAAAACCTCCCACACCTCCCCCTGAACCTGAAACATAAAATGAATGCAATTGTTGTTGTT

**Supplementary Figure S13**

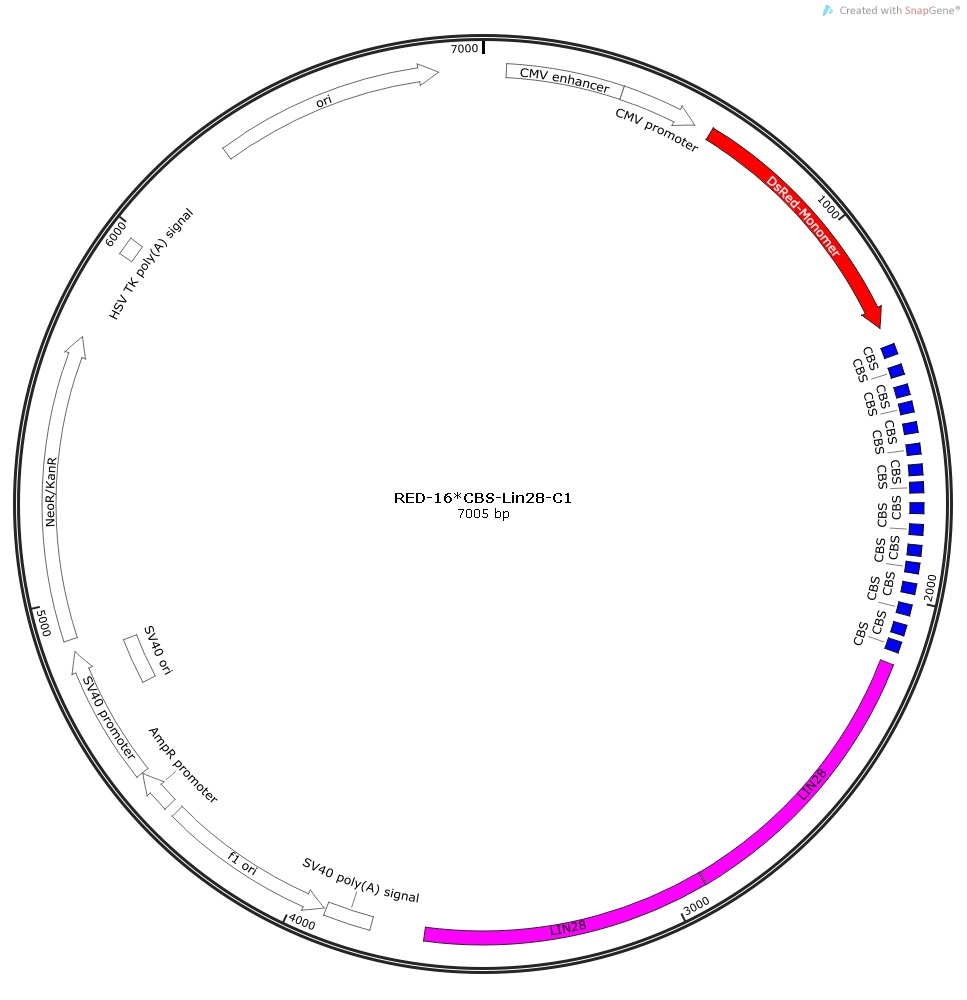

**Supplementary Figure S13.** RED-16×CBS-Lin28-C1 construct.

The nucleotide sequences of RFP, CBS, and Lin28 are marked in red, blue, and pink, respectively.

GGTTTAGTGAACCGTCAGATCCGCTAGCGCTACCGGTCGCCACCatgGACAACACCGAGGACGTCATCAAGGAGTTCATGCAGTTCAAGGTGCGCATGGAGGGCTCCGTGAACGGCCACTACTTCGAGATCGAGGGCGAGGGCGAGGGCAAGCCCTACGAGGGCACCCAGACCGCCAAGCTGCAGGTGACCAAGGGCGGCCCCCTGCCCTTCGCCTGGGACATCCTGTCCCCCCAGTTCCAGTACGGCTCCAAGGCCTACGTGAAGCACCCCGCCGACATCCCCGACTACATGAAGCTGTCCTTCCCCGAGGGCTTCACCTGGGAGCGCTCCATGAACTTCGAGGACGGCGGCGTGGTGGAGGTGCAGCAGGACTCCTCCCTGCAGGACGGCACCTTCATCTACAAGGTGAAGTTCAAGGGCGTGAACTTCCCCGCCGACGGCCCCGTAATGCAGAAGAAGACTGCCGGCTGGGAGCCCTCCACCGAGAAGCTGTACCCCCAGGACGGCGTGCTGAAGGGCGAGATCTCCCACGCCCTGAAGCTGAAGGACGGCGGCCACTACACCTGcGACTTCAAGACCGTGTACAAGGCCAAGAAGCCCGTGCAGCTGCCCGGCAACCACTACGTGGACTCCAAGCTGGACATCACCAACCACAACGAGGACTACACCGTGGTGGAGCAGTACGAGCACGCCGAGGCCCGCCACTCCGGCTCCCAGTCCGGACTCAGATCTCGACTTTGAGCGCTACCGGACTCAGATCTCGACCGAGTTCCCCGCGCCAGCGGGGATAAACCGCGCCAGCTTCGAATTCTGCAGTCGAAGAGTTCCCCGCGCCAGCGGGGATAAACCCGATCCACCGGTATTTATCTCGACCGAGTTCCCCGCGCCAGCGGGGATAAACCGCGCCAGCTTCGAATTCGAGTTCCCCGCGCCAGCGGGGATAAACCCGATCCACCGGTATTTATCTCGACCGAGTTCCCCGCGCCAGCGGGGATAAACCGCGCCAGCTTCGAATTCTGCAGTCGAAGAGTTCCCCGCGCCAGCGGGGATAAACCCGATCCACCGGTATTTATCTCGACCGAGTTCCCCGCGCCAGCGGGGATAAACCGCGCCAGCTTCGAATTCGAGTTCCCCGCGCCAGCGGGGATAAACCCGATCCACCGGTATTTATCTCGACCGAGTTCCCCGCGCCAGCGGGGATAAACCGCGCCAGCTTCGAATTCTGCAGTCGAAGAGTTCCCCGCGCCAGCGGGGATAAACCCGATCCACCGGTATTTATCTCGACCGAGTTCCCCGCGCCAGCGGGGATAAACCGCGCCAGCTTCGAATTCGAGTTCCCCGCGCCAGCGGGGATAAACCCGATCCACCGGTATTTATCTCGACCGAGTTCCCCGCGCCAGCGGGGATAAACCGCGCCAGCTTCGAATTCTGCAGTCGAAGAGTTCCCCGCGCCAGCGGGGATAAACCCGATCCACCGGTATTTATCTCGACCGAGTTCCCCGCGCCAGCGGGGATAAACCGCGCCAGCTTCGAATTCGAGTTCCCCGCGCCAGCGGGGATAAACCCGATCCACCGGTATTTATCTCGAGTTTGATAGGaaaccctccatcccttgttcccaacctcctaagtcaagaccattaccatttctttctttcttttttttttttttttaaaatggagtctcactgtgtcacccaggctggagtgcagtggcatgatcggctcactgcagcctctgcctcttgggttcaagtgattctcctgcctcagcctcctgagtagctgggatttcaggcacccgccacactcagctaatttttgtatttttagtagagacggggtttcaccatgttgtccaggctggtctggaactcctgacctcaggtgatctgcccaccttggcttcccaaagtgctgggattacaggcatgagccaccatgctgggccaaccatttcttggtgtattcatgccaaacacttaagacactgctgtagcccaggcgcggtggctcacacctgtaatcccagcactttggaaggctgaggcgggcggatcacaaggtcacgagttcaaaactatcctggccaacacagtgaaaccccgtctctactaaaatacaaaaaaattagccgggtgtggtggtgcatgcctttagtcctagctattcaggaggctgaggcaggggaatcgcttgaacccgagaggcagaggttgcagtgagctgagatcgcaccactgcactccagcctggttacagagcaagactctgtctcaaacaaaacaaaacaaaacaaaaacacactactgtattttggatggatcaaacctccttaaCCGaaaccctccatcccttgttcccaacctcctaagtcaagaccattaccatttctttctttcttttttttttttttttaaaatggagtctcactgtgtcacccaggctggagtgcagtggcatgatcggctcactgcagcctctgcctcttgggttcaagtgattctcctgcctcagcctcctgagtagctgggatttcaggcacccgccacactcagctaatttttgtatttttagtagagacggggtttcaccatgttgtccaggctggtctggaactcctgacctcaggtgatctgcccaccttggcttcccaaagtgctgggattacaggcatgagccaccatgctgggccaaccatttcttggtgtattcatgccaaacacttaagacactgctgtagcccaggcgcggtggctcacacctgtaatcccagcactttggaaggctgaggcgggcggatcacaaggtcacgagttcaaaactatcctggccaacacagtgaaaccccgtctctactaaaatacaaaaaaattagccgggtgtggtggtgcatgcctttagtcctagctattcaggaggctgaggcaggggaatcgcttgaacccgagaggcagaggttgcagtgagctgagatcgcaccactgcactccagcctggttacagagcaagactctgtctcaaacaaaacaaaacaaaacaaaaacacactactgtattttggatggatcaaacctccttaaAAGCTTGGGGGGATCCACCGGATCTAGATAACTGATCATAATCAGCCATACCACATTTGTAGAGGTTTTACTTGCTTTAAAAAACCTCCCACACCTCCCCCTGAACCTGAAACATAAAATGAATGCAATTGTTGTTGTT

**Supplementary Figure S14**

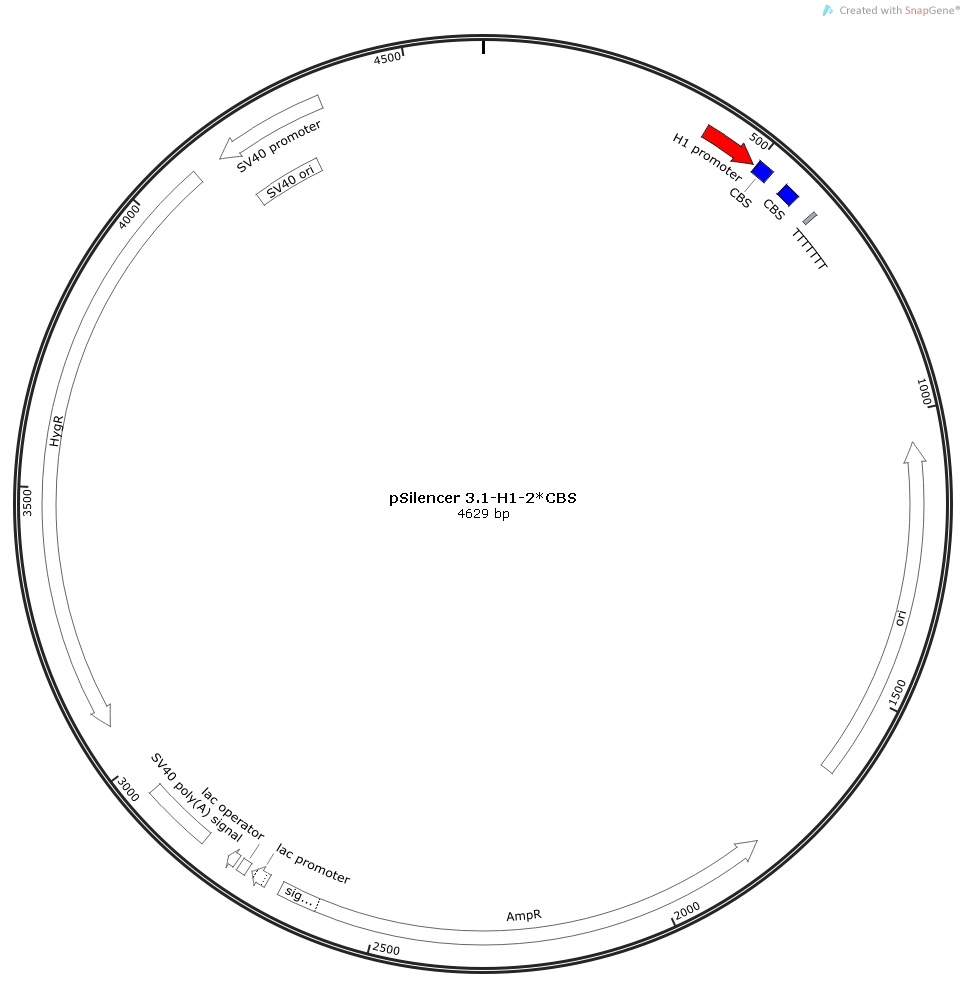

**Supplementary Figure S14.** pSilencer 3.1-H1-2×CBS construct.

The nucleotide sequences of H1 promoter and CBS are marked in red and blue, respectively.

GAATTCATATTTGCATGTCGCTATGTGTTCTGGGAAATCACCATAAACGTGAAATGTCTTTGGATTTGGGAATCTTATAAGTTCTGTATGAGACCACTCGGATCCgagttccccgcgccagcggggataaaccgCTACTACCTCATTGATCCTGGCTTGCTAGCgagttccccgcgccagcggggataaaccgAGATCTaaGGTACCacGAGCTCtgCTCGAGaAGCGCTTTTTTTTAAGCTTGGCGTA

**Supplementary Figure S15**

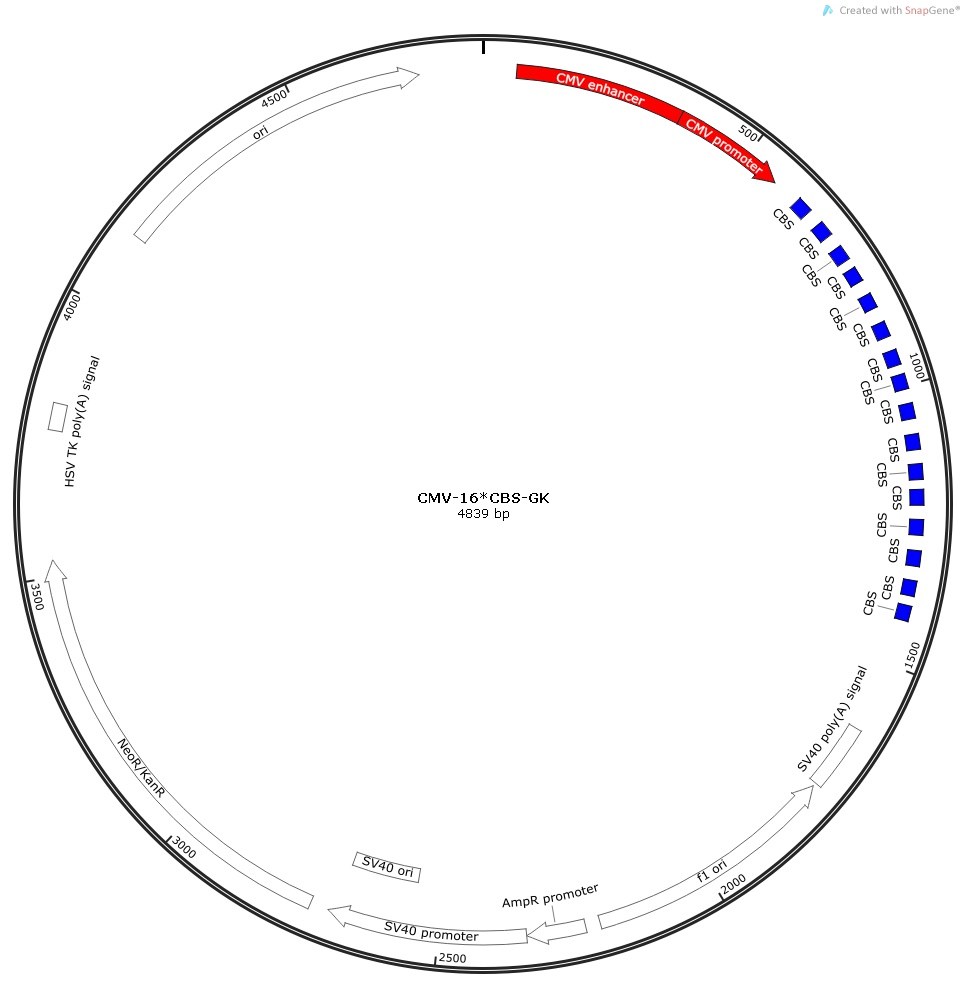

**Supplementary Figure S15.** CMV-16×CBS-GK construct.

The nucleotide sequences of CMV promoter and CBS are marked in red and blue, respectively.

CGTTACATAACTTACGGTAAATGGCCCGCCTGGCTGACCGCCCAACGACCCCCGCCCATTGACGTCAATAATGACGTATGTTCCCATAGTAACGCCAATAGGGACTTTCCATTGACGTCAATGGGTGGAGTATTTACGGTAAACTGCCCACTTGGCAGTACATCAAGTGTATCATATGCCAAGTACGCCCCCTATTGACGTCAATGACGGTAAATGGCCCGCCTGGCATTATGCCCAGTACATGACCTTATGGGACTTTCCTACTTGGCAGTACATCTACGTATTAGTCATCGCTATTACCATGGTGATGCGGTTTTGGCAGTACATCAATGGGCGTGGATAGCGGTTTGACTCACGGGGATTTCCAAGTCTCCACCCCATTGACGTCAATGGGAGTTTGTTTTGGCACCAAAATCAACGGGACTTTCCAAAATGTCGTAACAACTCCGCCCCATTGACGCAAATGGGCGGTAGGCGTGTACGGTGGGAGGTCTATATAAGCAGAGCTGGTTTAGTGAACCGTCAGATCCGCTAGCGCTACCGGACTCAGATCTCGACCGAGTTCCCCGCGCCAGCGGGGATAAACCGCGCCAGCTTCGAATTCTGCAGTCGAAGAGTTCCCCGCGCCAGCGGGGATAAACCCGATCCACCGGTATTTATCTCGACCGAGTTCCCCGCGCCAGCGGGGATAAACCGCGCCAGCTTCGAATTCGAGTTCCCCGCGCCAGCGGGGATAAACCCGATCCACCGGTATTTATCTCGACCGAGTTCCCCGCGCCAGCGGGGATAAACCGCGCCAGCTTCGAATTCTGCAGTCGAAGAGTTCCCCGCGCCAGCGGGGATAAACCCGATCCACCGGTATTTATCTCGACCGAGTTCCCCGCGCCAGCGGGGATAAACCGCGCCAGCTTCGAATTCGAGTTCCCCGCGCCAGCGGGGATAAACCCGATCCACCGGTATTTATCTCGACCGAGTTCCCCGCGCCAGCGGGGATAAACCGCGCCAGCTTCGAATTCTGCAGTCGAAGAGTTCCCCGCGCCAGCGGGGATAAACCCGATCCACCGGTATTTATCTCGACCGAGTTCCCCGCGCCAGCGGGGATAAACCGCGCCAGCTTCGAATTCGAGTTCCCCGCGCCAGCGGGGATAAACCCGATCCACCGGTATTTATCTCGACCGAGTTCCCCGCGCCAGCGGGGATAAACCGCGCCAGCTTCGAATTCTGCAGTCGAAGAGTTCCCCGCGCCAGCGGGGATAAACCCGATCCACCGGTATTTATCTCGACCGAGTTCCCCGCGCCAGCGGGGATAAACCGCGCCAGCTTCGAATTCGAGTTCCCCGCGCCAGCGGGGATAAACCCGATCCACCGGTATTTATCTCGAGCTCAAGCTTCGAATTCTGCAGTCGACGGTACCGCGGGCCCGGGATCCATATGTAAGTAAGTAAGCGGCCGCGACTCTAGATCATAATCAGCCATACCACATTTGTAGAGGTTTTACTTGCTTTAAAAAACCTCCCACACCTCCCCCTGAACCTGAAACATAAAATGAATGCAATTGTTGTTGTT

**Supplementary Figure S16**

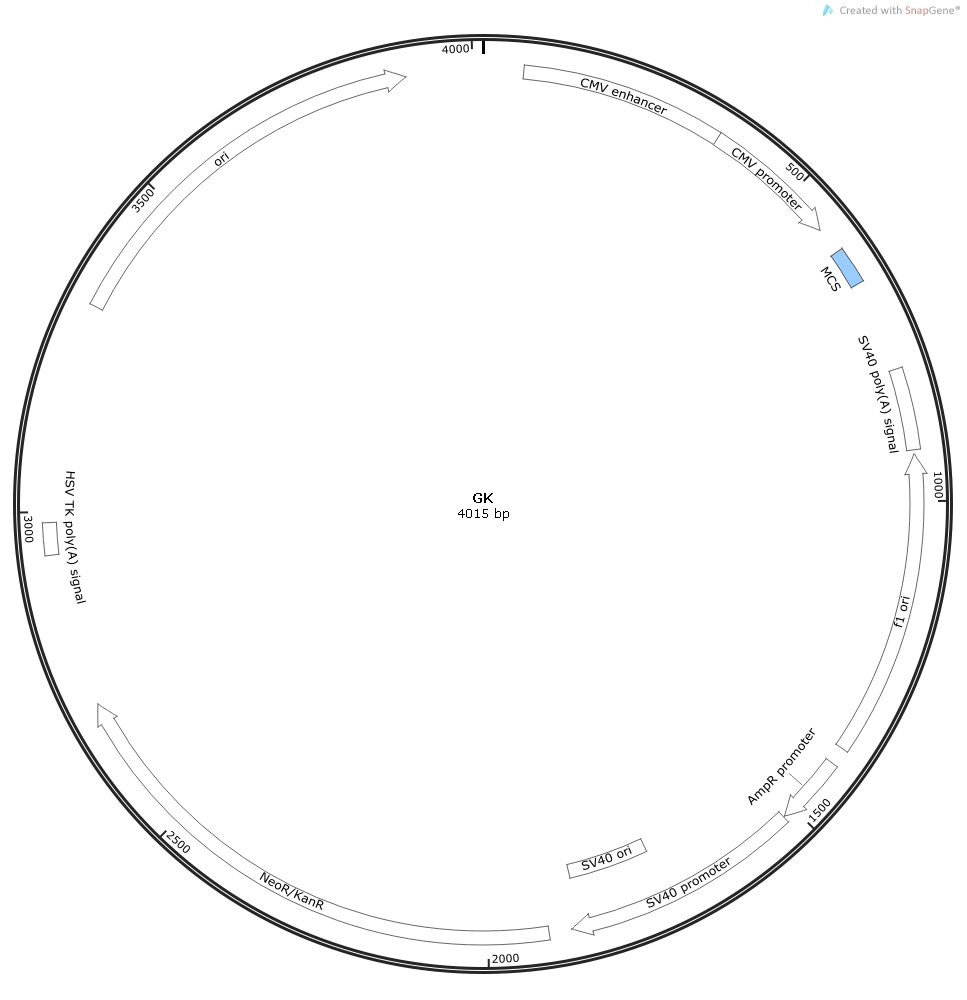

**Supplementary Figure S16.** GK construct.

The nucleotide sequence of multiple cloning site (MCS) is marked in blue.

GGTTTAGTGAACCGTCAGATCCGCTAGCGCTACCGGACTCAGATCTCGAGCTCAAGCTTCGAATTCTGCAGTCGACGGTACCGCGGGCCCGGGATCCATATGTAAGTAAGTAAGCGGCCGCGACTCTAGATCATAATCAGCCATACCACATTTGTAGAGGTTTTACTTGCTTTAAAAAACCTCCCACACCTCCCCCTGAACCTGAAACATAAAATGAATGCAATTGTTGTTGTT
